# Decoding MASLD Progression: A Molecular Trajectory-Based Framework for Modelling Disease Dynamics

**DOI:** 10.1101/2025.01.14.632908

**Authors:** Ioannis Kamzolas, Thodoris Koutsandreas, Charlie George Barker, Anna Vathrakokoili Pournara, Harry Weston, Naoto Fujiwara, Yujin Hoshida, Quentin M. Anstee, Michele Vacca, Irene Papatheodorou, Antonio Vidal-Puig, Evangelia Petsalaki

## Abstract

Metabolic Dysfunction-Associated Steatotic Liver Disease (MASLD) has emerged as a silent pandemic, affecting nearly one-third of the global population. MASLD encompasses a spectrum of liver disorders, ranging from simple steatosis to Metabolic Dysfunction-Associated Steatohepatitis (MASH), characterised by lipotoxicity, hepatocellular injury, inflammation, and fibrosis, which can eventually progress to cirrhosis and hepatocellular carcinoma. Despite the progressive nature of MASLD/MASH, current research and clinical practice primarily rely on static, histopathology-defined stages that fail to capture the continuous nature of disease progression.

Here, we present an integrative framework that combines patient pseudo-temporal ordering, network analysis, and cell-type deconvolution to reconstruct the continuous MASLD/MASH trajectory. By analysing patient liver transcriptomic profiles, we position patients along this data-driven trajectory, moving beyond conventional stage-based classifications. This approach reveals the sequence of critical molecular events underlying MASLD/MASH progression, providing mechanistic insights into the disease’s pathophysiology. By integrating these findings with plasma proteomics data, we identify novel trajectory-specific plasma biomarkers that predict disease stage (and trajectory position) independently of histology.

Together, these findings demonstrate the value of trajectory-based frameworks for understanding MASLD pathophysiology and highlight new opportunities for precision diagnosis and therapeutic target prioritisation across the disease spectrum.

## Introduction

Metabolic Dysfunction-Associated Steatotic Liver Disease (MASLD) encompasses a spectrum of conditions, ranging from simple steatosis to Metabolic Dysfunction-Associated Steatohepatitis (MASH), which can progress to fibrosis, cirrhosis, and hepatocellular carcinoma^1^. This global epidemic affects nearly one-third of Western populations and is primarily driven by the rising prevalence of metabolic syndrome, type 2 diabetes, and obesity^2,3^. Despite its prevalence, most MASLD cases remain asymptomatic and undiagnosed until advanced stages.

Diagnosis of MASLD relies on imaging (ultrasound, MRI spectroscopy) and non-invasive fibrosis scores such as the NAFLD Fibrosis Score and FIB-4, developed initially to detect advanced fibrosis^4,5^. Although liver biopsy remains the histological gold standard, including for derivation of the NAFLD Activity Score^6^, its use is limited by sampling variability and procedural risk^7,8^. Current AASLD and EASL–EASD–EASO guidelines endorse non-invasive tests but emphasise the need for complementary diagnostic strategies^9,10^. Elastography methods, including transient elastography (FibroScan)^11^ and magnetic resonance elastography, have improved fibrosis assessment, but are limited by precision, accessibility, or cost. Consequently, despite improved histological scoring reproducibility^12^, robust non-invasive biomarkers for MASLD remain lacking.

Advances in systems biology and multi-omics have accelerated MASLD/MASH research^13–17^, with transcriptomic, proteomic, and metabolomic studies linking the disease to metabolic syndrome, insulin resistance, and processes such as lipid accumulation, ER stress, and oxidative stress^18^. Genome-wide association studies (GWAS) have identified major genetic risk factors, most notably the PNPLA3 I148M variant^19–21,22^, which was later validated across large multi-ethnic cohorts^23^. Additional variants in TM6SF2^24^, MBOAT7^25^, and HSD17B13^26^ further define the genetic landscape, implicating lipid metabolism, inflammation, and fibrosis pathways. Despite these breakthroughs, the complexity of MASLD progression, shaped by genetic, metabolic, and environmental interactions, demands further research to elucidate stage-specific mechanisms and distinguish benign cases from progressive disease.

Histology has long been a cornerstone for understanding MASLD pathophysiology, but its limitations as a sole proxy for disease mechanisms are increasingly evident. Pseudo-temporal ordering, initially developed for microarray data^27^ and now widely applied in single-cell genomics, provides an alternative framework by arranging cells or patients along trajectories based on their molecular states. This approach has recently been used to study Alzheimer’s disease progression through brain expression data^28,29^, and offers a promising, histology-independent method for investigating MASLD progression via transcriptomic profiles. By complementing traditional assessments, pseudo-temporal ordering provides a data-driven perspective on the molecular changes underpinning disease progression.

In this study, we applied pseudo-temporal ordering to human MASLD/MASH liver transcriptomes and integrated the resulting progression axis with co-expression and regulatory network analyses. This framework resolved dynamic, stage-dependent transcriptional programmes across the disease continuum and was reproducible across independent cohorts, including longitudinal paired biopsies. By intersecting the disease network with liver–plasma-correlated proteins, we further derived a 57-gene plasma-accessible biomarker panel that outperformed established non-invasive scores for advanced fibrosis and enabled continuous patient positioning along the molecular trajectory. Together, these results provide a scalable strategy for mapping MASLD progression from cross-sectional data and for supporting non-invasive staging and longitudinal monitoring.

## Results

### Transcriptomics-based disease trajectory analysis captures MASLD progression

We hypothesised that MASLD progression could be modelled as a continuum of patient states. Given variability between patients, including the role of sex^30^, comorbid conditions^31^ and other parameters in the timing of disease progression^32^, we focused on modelling the molecular underpinnings of the progression that are directly related to the histological phenotypes, as these are common to all patients.

To test this hypothesis, we analysed RNA-seq data from 136 patients across two published cohorts^14,33^, adjusting for sex during data integration (Methods; **Supplementary Tables 1 & 2**). After excluding one outlier, we applied pseudo-temporal ordering to derive a transcriptomics-based disease trajectory (**Fig. 1A–B**; **Supplementary Fig. 1A-B**). The inferred trajectory showed strong concordance with histological measures, including steatosis, ballooning, inflammation, fibrosis, and NAS (Pearson R = 0.96–1.0; p< 0.05 for most compared disease stages; ANOVA followed by Tukey’s pairwise comparisons; **Supplementary Fig. 1C**), confirming alignment with established disease stages.

**Figure 1.**
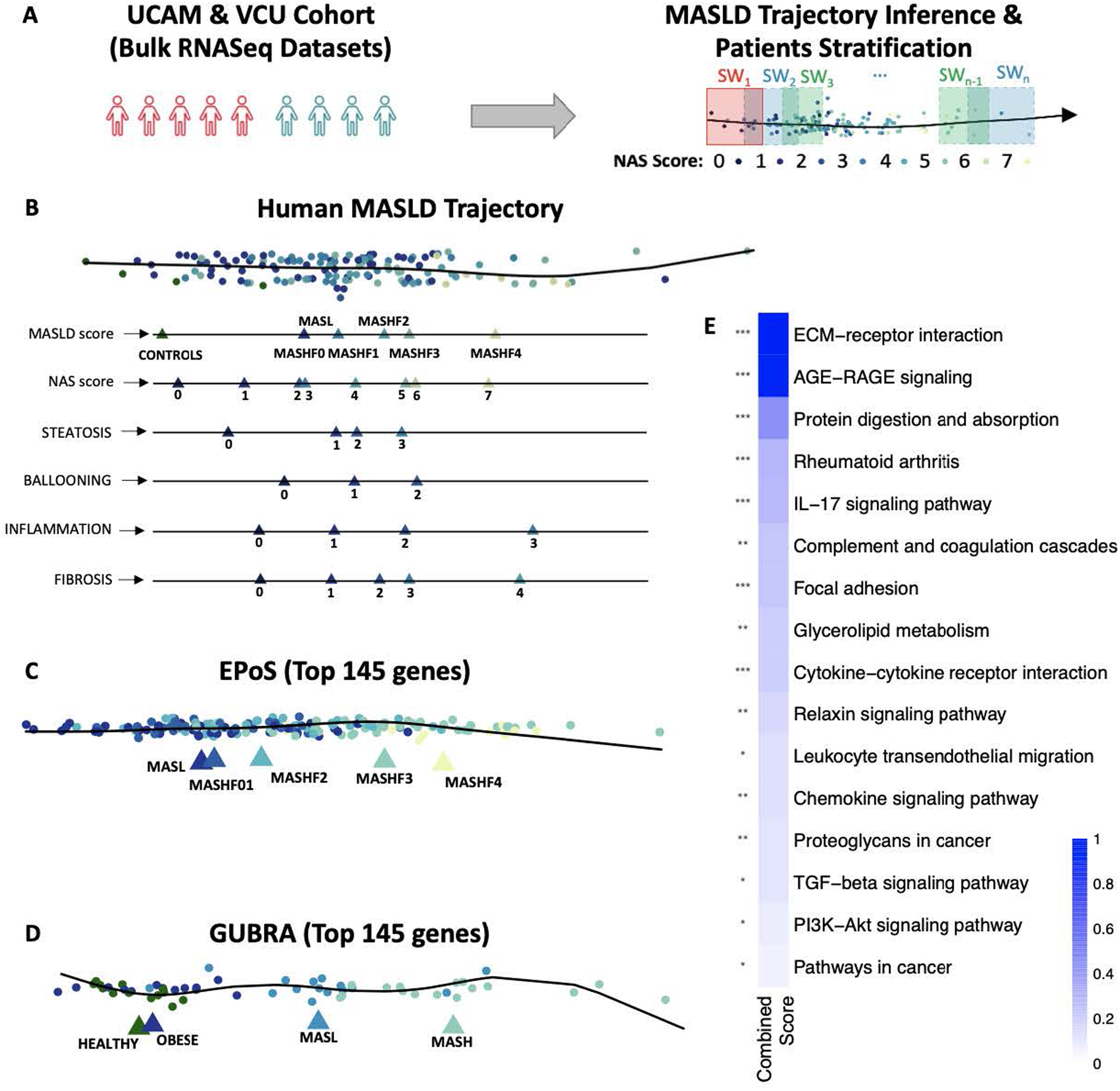
Pseudo-temporal ordering of patients captures disease progression and identifies MASLD gene signature. **A.** Schematic of pseudotemporal ordering of patients based on bulk transcriptomics data from liver biopsies and stratification of patients into sliding windows. **B.** Pseudo-temporal ordering of patients based on transcriptomic data recapitulates disease progression based on individual phenotypes, i.e., steatosis, ballooning, inflammation and fibrosis, and based on the NAS and MASLD scores. **C-D.** Pseudotemporal ordering of two independent orthogonal datasets, based on the 145 genes that are most predictive of the trajectory, provides a linear and clear separation of the disease stages (EPoS dataset (**C**) and Gubra dataset (**D**)). Triangles show the average position of each histopathologically characterised stage on the trajectory. **E.** Functional enrichment analysis results of these 145 genes using EnrichR, where the Combined Score (c) has been calculated as c = log(p_value) * z_score, with p_value representing the Fisher exact test outcome, and z-score the deviation from the expected rank.

To validate the trajectory model, we tested it on two independent RNA-Seq datasets: the EPoS dataset, a large multi-cohort dataset encompassing 168 MASLD patients across the full disease spectrum, and the Gubra dataset, comprising 26 healthy normal weight and obese individuals, and 31 MASLD and MASH patients^13,34^. Initial pseudotemporal ordering of these patients showed a trend of disease progression; however, additional variance in both datasets was observed that did not fully correlate with disease progression (**Supplementary Fig. 2A-B**).

To identify the principal drivers of variance along the MASLD/MASH trajectory and assess their generalisability, we applied a random forest approach to the discovery cohorts (UCAM/SANYAL), identifying 145 genes predictive of disease stage and histopathological features (AUC 0.62–0.73; **Supplementary Table 3**). This gene set more accurately recapitulated the disease trajectory in independent datasets^13,34^, improving linearity and stage separation compared with the full transcriptome (**Fig. 1C–D**). Feature selection was performed exclusively on the discovery data, ensuring no information leakage and supporting the model’s robustness across the larger, more heterogeneous EPoS and Gubra cohorts.

Finally, we evaluated the ability of our 145 gene signature to place longitudinal data from 58 patients from a different ethnic background (Japanese cohort^35^) along the trajectory. We found that their location on the trajectory was largely consistent with the changes in their histological profile, in particular where their disease regressed (**Supplementary Fig. 2C-E**). Together, these results demonstrate the generalisability of our 145 genes for MASLD trajectory inference.

The identified genes were enriched in pathways central to MASLD/MASH progression, including ECM-receptor interaction and focal adhesion (associated with fibrosis), AGE-RAGE and IL-17 signalling (inflammatory responses), and TGF-beta and PI3K-Akt signalling (wound healing and metabolic dysregulation). Other pathways, such as glycerolipid metabolism and cytokine-receptor interaction, reflect the metabolic and immune dysregulation characteristic of MASLD/MASH (**Fig. 1E**, **Supplementary Table 4**).

To enable deeper insights into the molecular changes driving MASLD progression, we next combined a sliding window (SW) approach, dividing patients into groups along the disease trajectory, with network analysis. To optimise the sliding windows sequence, we developed a graph-based method that maximised the information content of each window (Methods). Using this approach, patients were divided into 13 subgroups along the trajectory for functional network analysis (Methods; **Fig. 1A**; **Supplementary Fig. 3**; **Supplementary Tables 1 & 5**).

### Data-Driven Global MASLD/MASH Network Recapitulates Key Molecular Mechanisms of Disease

First, to construct a MASLD reference network, we adapted our previously published method for generating phenotype-specific networks by integrating paired transcriptomics and phenotype data^36^ (**Fig. 2A**; Methods). Using Weighted Gene Co-expression Network Analysis (WGCNA)^37^, we identified gene co-expression modules and linked them to key phenotypic features, including the NAS score, steatosis, ballooning, and fibrosis (**Fig. 2B**; **Supplementary Table 6**). The ballooning score reflects hepatocyte injury, whereas the inflammation score, which reflects lobular immune infiltrates. The inflammation score was used as a covariate in the analysis, due to difficulty deconvolving its role as a cause vs. effect and because it dominated the signal. Nonetheless it is already represented in the NAS score. Finally, to extract the modules associated with our histological phenotypes of interest, we used linear regression, identifying 10 modules associated with at least one phenotypic feature (**Fig. 2B**; Methods). After filtering genes that were not correlated with the eigengene of each module (Methods), the number of genes in each significant module ranged from 192 in the MEsalmon to 2949 in the MEturquoise module (**Supplementary Table 6**).

**Figure 2.**
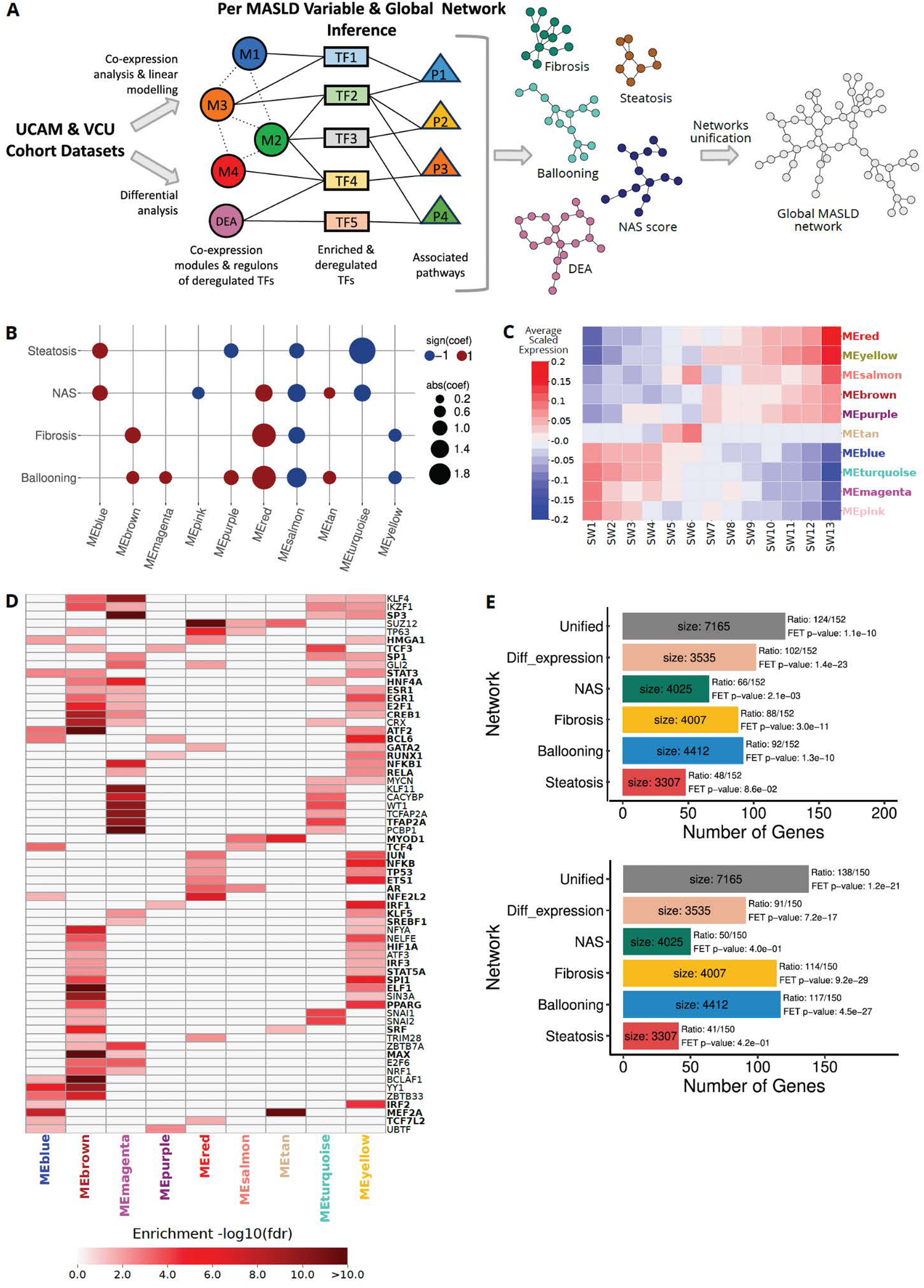
Phenotype-specific gene modules associated with the different MASLD variables. **A.** Schematic of the approach to extract a reference MASLD network. **B.** Association (coefficient from linear model) of the different gene co-expression modules with NAS score, Steatosis, Ballooning, and Fibrosis. Red indicates a positive coefficient, whereas blue indicates negative coefficient. **C.** Heatmap is showing the average scaled expression of the modules along the pseudotemporal trajectory. The values for each module were calculated by averaging the scores of SW-grouped samples along its first principal component (eigengene). **D.** TFs whose regulons are enriched in the modules. The figure shows TFs per module, coloured according to the adjusted p-value of the enrichment test. The presented TFs have been identified as enriched in multiple modules. Bold indicates TFs identified as deregulated (FDR<0.05) in at least one sliding window (see sliding window analysis results). The complete of TFs derived from the enrichment analysis is given in **Supplementary Table 8. E.** Enrichment of MASLD network and its components in known MASLD genes curated from the literature (top) or from MSigDB (bottom).

Reactome^38^ enrichment of the phenotype-associated modules highlighted patterns consistent with MASLD progression (**Supplementary Table 7**). MEbrown, positively associated with ballooning and fibrosis, showed exceptionally strong enrichment for protein translation (R-HSA-72766, FDR = 1.9e-99) and ribosome biogenesis (R-HSA-72706, FDR = 5.5e-59), reflecting hepatocyte stress and increased biosynthetic demand during injury^39^. MEred, also linked to fibrosis and ballooning, was enriched for extracellular matrix organization (R-HSA-1474244, FDR = 6.5e-23), aligning with well-established fibrogenic remodeling^40^. In contrast, MEblue, associated with steatosis and NAS, was enriched for gene-regulatory programs (R-HSA-74160, FDR = 1.6e-18) consistent with early metabolic and transcriptional reprogramming in lipid-laden hepatocytes^41^. Finally, MEyellow, although negatively associated with fibrosis and ballooning, showed strong enrichment for immune system pathways (R-HSA-168256, *FDR = 1.8e-47*), capturing the inflammatory processes characteristic of advanced disease. Together, these phenotype–pathway correspondences provide confidence that the reference network recapitulates key molecular processes underlying steatosis, hepatocellular injury, inflammation, and fibrosis in MASLD.

As a confirmation that our modules are indeed following the expected dynamics reflected in the histology, we evaluated the ‘activity’ of the modules along our sliding window identifying two groups (Methods): Modules that are generally associated with Fibrosis, Ballooning and NAS (e.g. MEred, MEbrown and others; **Fig 2B-C**) being less active in the beginning of the trajectory and peaking at the later stages, whereas those associated with Steatosis and NAS (e.g. MEblue, MEturqoise) behaving oppositely. Interestingly, METan, which is mainly associated with ballooning and NAS, only shows activity in the early-mid stages (SW5-7), in agreement with the histology.

Transcription factor (TF) enrichment analysis of the phenotype-associated modules identified 199 TF regulons across nine out of the ten significant modules (FDR < 0.05), with many converging on TGF-beta-driven fibrogenic signalling (**Supplementary Table 8**). Among these, 65 TFs were shared across multiple modules (**Fig. 2D**). Core TGF-beta regulators SMAD2, 3 and 4, were selectively enriched in the MEred module (FDR = 0.04, 5e-05, 6.3e-08 respectively; **Supplementary Table 8**), consistent with its link to ballooning and fibrosis (**Fig. 2B**). Enriched in the ballooning and fibrosis-associated modules, we found SP1, SRF, and ETS1, known to interact with TGF-beta signalling to promote the expression of tissue remodelling and fibrosis factors^42–44^. EGR1, a TGF-beta–responsive TF previously linked to both steatosis and fibrosis^45,46,47^, was enriched across multiple modules, bridging early and late disease features (**Fig. 2D**).

HNF4A, a key regulator of liver development and morphogenesis^48^, and PPARG, which controls lipid storage and adipocyte differentiation in liver^49^, appeared across modules with opposing phenotype associations (MEbrown versus MEturquoise or MEyellow; FDR HNF4A = ∼0.001, PPARG = 1e-04 and 4.3e-05; **Fig. 2B & D**), indicating divergent transcriptional programmes across disease stages. SREBF1, found in MEyellow (FDR = 0.05) and MEmagenta (FDR = 0.04), similarly bridges lipogenic regulation and early stress responses, consistent with its shift from metabolic control toward activation under hepatocellular injury^33,50^.

Inflammatory and hypoxia-responsive TFs (NFKB1, RELA, HIF1A) were enriched in MEyellow (FDR = 6.8e-11, 9.6e-11, and 0.006 respectively), whereas CREB1 showed strongest enrichment in MEbrown (FDR = 2.0e-24), reflecting distinct immune-driven^51,52,53^ versus hepatocyte-intrinsic^54^ stress transcriptional contexts (**Fig. 2B & D; Supplementary Table 8**).

To ensure comprehensive regulatory coverage, we additionally incorporated TFs differentially regulated along the disease trajectory (FDR < 0.05; Methods), yielding a combined TF module (DEA), in which 85 of the 199 enriched TFs showed stage-specific deregulation. (**Supplementary Table 8**).

We then integrated the phenotype-associated modules, their enriched TFs, and corresponding pathways (**Supplementary Table 9**; Methods) to construct module-specific regulatory networks for each histological feature, as well as for our dynamically regulated TFs along the disease trajectory (**Fig. 2A**). These networks were then merged into a comprehensive MASLD/MASH disease network, henceforth referred to as the MASLD network for simplicity, to capture dynamic changes in cellular processes across disease stages (**Fig. 2A**; Methods; **Supplementary Table 10**). The resulting network contained 7,165 nodes and showed significant enrichment for previously reported MASLD-associated genes from both our curated literature set (**Supplementary Table 11**) and an established MSigDB^55^ gene set (MSigDB ID: M39806; Methods) (**Fig. 2E**).

As an orthogonal validation we used the same pipeline to generate a MASLD network using the larger EPoS dataset described above^13,34^. That network comprised 6,732 nodes (**Supplementary Table 10**) and the intersection of the two networks was 5,241 nodes (Jaccard index=0.61, odds ratio=7.11, Fisher’s exact test p-value=0).

### Sliding Window Analysis Highlights Molecular Dysregulation in MASLD Progression

We next sought to combine our reference MASLD network with our patient trajectory by using our sliding window approach (**Fig 1A**) to explore the molecular mechanisms unerlying MASLD progression relevant to observed histological phenotypes.

Initially, we identified TFs that were dynamically regulated across MASLD progression, showing strong concordance with traditional stage-based stratifications (early, middle, late and NAS score-based patient stratification), while providing greater temporal resolution (**Fig. 3A**; **Supplementary Table 12**). In total, 122 TFs exhibited at least one significant deregulation event along the trajectory (FDR < 0.05), of which 57 were also detected by early–middle–late or NAS-based analyses with concordant directionality. This shared set included TF clusters enriched for Toll-like receptor signalling (e.g. FOS, JUN, CREB1, TP53, NFKB1/2, RELA; R-HSA-168898, FDR = 5.3e-05), estrogen receptor–mediated signalling (e.g. ESR1, RUNX1, MYB; R-HSA-8939211, FDR = 6.4e-07), and cytokine signalling (e.g. EGR1, IRF and STAT family members; R-HSA-1280215, FDR = 2.8e-08) (**Supplementary Figure 4**).

**Figure 3.**
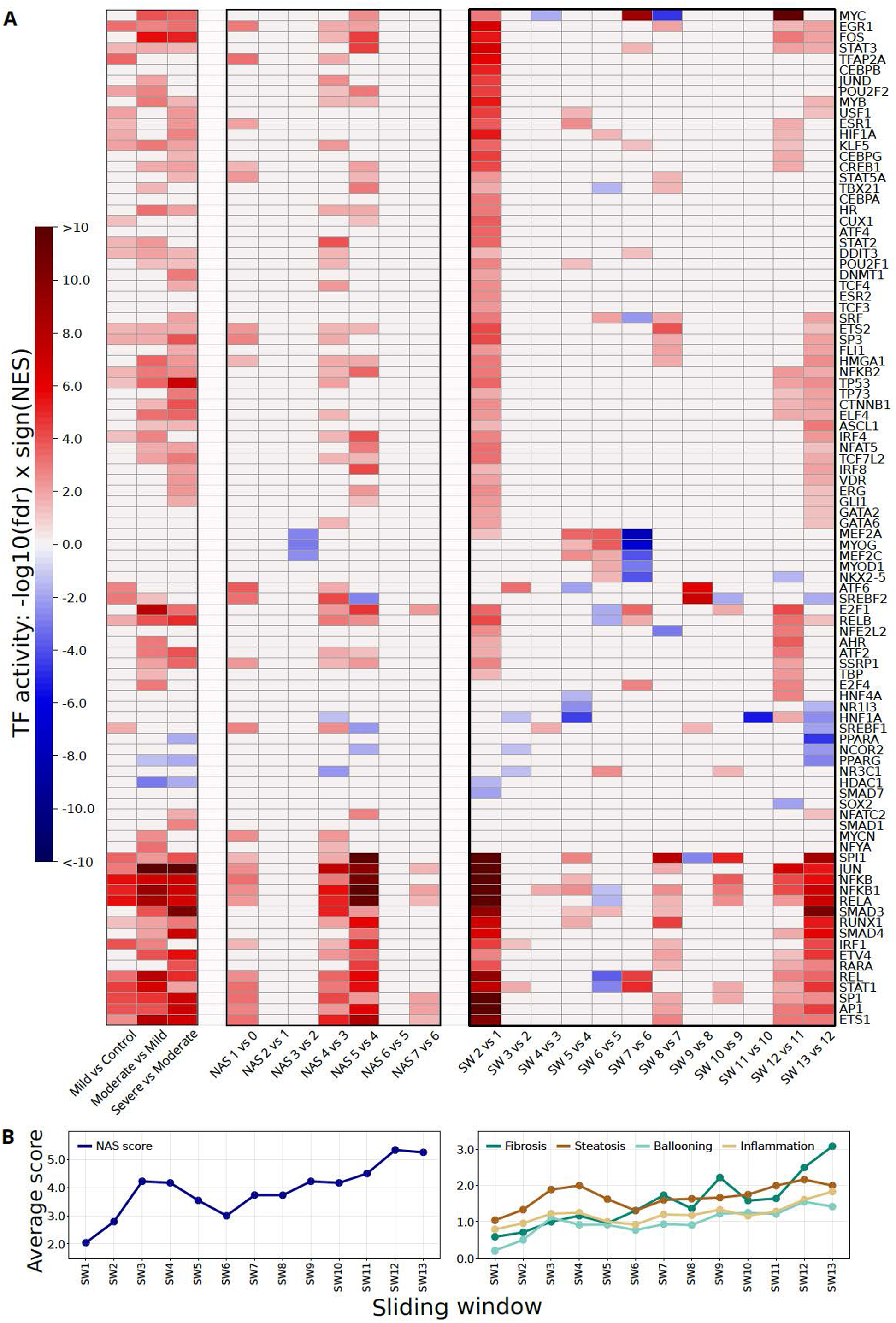
Sliding window analysis to study the landscape of molecular changes along the MASLD trajectory. **A.** Results from TF activity analysis in two types of discrete patient stratification (Mild, Moderate, Severe and NAS scores), and the pseudotemporal trajectory. The colour indicates a change in TF activity compared to the previous disease stage. **B.** Mean phenotype and NAS pathologist scores along the SW trajectory.

SREBF1 was consistently upregulated in early and intermediate disease states across all stratifications (in SW4 and SW9 according to the SW approach), confirming its pro-steatotic role^33,56^, but only the trajectory-based and NAS-based approaches captured its downregulation in later stages^57^, illustrating the benefit of increased granularity.

Beyond this shared signal, the trajectory-based approach uniquely identified several key regulatory events missed by discrete classifications, including early downregulation of HNF4A (SW5), NR1I3 (SW5), and HNF1A (SW3 & 5), reflecting disrupted hepatocyte metabolic identity, inflammatory signalling, and tissue remodelling^58,59,60^ (**Fig. 3A**). Conversely, sustained upregulation of pro-fibrotic regulators such as SRF^43^ and dynamic regulation of TGF-β pathway components, including downregulation of the inhibitory SMAD7^61^ (SW2), were detected only along the SW trajectory (**Fig. 3A**). Additional TFs (e.g. MYC, STAT1, NFKB1) displayed complex, stage-dependent regulation, underscoring the ability of trajectory-based analysis to capture nuanced and biologically meaningful regulatory dynamics during MASLD progression (**Fig. 3A**).

These results were largely validated in an orthogonal analysis of the EPoS dataset^13,34^, where 69 TFs were deregulated in at least one SW, 57 of which overlapped with our primary analysis (Jaccard index = 0.43, odds ratio = 76.03, Fisher’s exact test p-value < 2.2e-16). Deregulation directionality, based on cumulative activation scores, showed high concordance between datasets with an average Pearson correlation coefficient (PCC) of 0.56 (**Supplementary figure 5A**). The average PCC of cumulative activities per SW was 0.38, revealing two opposing trends along the disease trajectory (**Supplementary figure 5B**). Notably, PCC decreased monotonically in early stages (SW2–SW5), indicating divergence in early molecular profiles between cohorts, likely due to the absence of healthy samples and higher MASLD variable values in SW1 of the EPoS dataset,which makes this group more similar to SW2 than in the UCAM/VCU cohort. In contrast, PCC increased in middle and late stages, demonstrating that despite cohort-specific differences, the trajectory-based approach captures consistent molecular programmes as disease progresses (**Supplementary figure 5B**).

An intriguing cluster of TFs, including MEF2A, MEF2C, MEF2D, MYOG, NKX2-5, and ZBTB4, exhibited unique activity patterns in our trajectory analysis that were not detected using traditional discrete patient staging (**Fig. 3A**). Their activity peaked during SW5–SW6 and then sharply declined at SW7, suggesting either a resolution of activation or a shift in regulatory dynamics. Notably, no significant changes in their activity were observed in later stages, indicating a potential stabilisation of their regulatory influence as the disease progressed (**Fig 3A**). These TFs, known primarily for their roles in myogenesis^62^, have also been implicated in hepatic stellate cell activation and their transition to a myofibroblast-like phenotype, a critical driver of fibrosis^63^. MEF2A/MEF2C and NKX2-5 also have documented roles in macrophage differentiation and inflammatory programming^64–66^, although not specifically related to MASLD. Further research is needed to understand the role of this TF cluster in MASLD progression.

To explore the underlying molecular processes, we employed a network propagation-based strategy to extract differentiated network signatures for each sliding window from the MASLD reference network (Methods). Reactome analysis of these networks identified 111 pathways exhibiting progressive changes along the disease trajectory (**Supplementary Fig. 6**; **Supplementary Table 13; Methods**). Signal transduction, including TGF beta, Receptor Tyrosine Kinase and others, and multiple immune pathways increased with disease progression (**Supplementary Fig. 6**), consistent with escalating inflammatory and signalling dysregulation. Extracellular matrix organisation pathways, linked to fibrosis, such as integrin and non-integrin membrane-ECM interactions, were upregulated predominantly from mid to late stages (**Supplementary Fig. 6**). Metabolic pathways showed more complex dynamics: lipid metabolism was initially downregulated (SW3-5) but showed modest upregulation after SW8, whereas glucose metabolism increased persistently from early stages, consistent with insulin resistance^67^. Orthogonal validation in the EPoS dataset^13,34^ identified 95 associated pathways, with 72 overlapping (Jaccard index = 0.54, odds ratio = 7.23, Fisher’s exact test p-value = 6.3e-12; **Supplementary Table 13**). Pathway deregulation directionality was concordant across datasets (average PCC = 0.46; **Supplementary Fig. 7A**), and cumulative pathway activities per SW showed even higher agreement (average PCC = 0.52; **Supplementary Fig. 7B**), mirroring oscillatory patterns observed in TF-based analyses but with higher overall correlation.

Finally, we compared pathway dysregulation across sliding windows with histopathological scores (steatosis, ballooning, inflammation, fibrosis, and NAS; **Fig. 3B**). Although these scores increased overall with disease progression, several pathways showed dysregulation earlier than histological changes. For example, extracellular matrix pathways were upregulated at SW8, receding the overall increase in fibrosis observed later along the trajectory., aligned with significant changes in multiple TFs (**Fig. 3A**) in that stage, indicating that molecular readouts may detect MASLD progression earlier than conventional assessments.

Overall, the enhanced resolution of our sliding window-based approach provides a comprehensive molecular landscape of MASLD progression, shedding light on the interplay between immune activation, fibrotic remodelling, and metabolic dysregulation over time.

### Cell-Type Deconvolution Along the MASLD Trajectory Reveals Network Changes Associated with Tissue Composition

Changes in liver cell composition and state are hallmarks of MASLD progression and reflect injury-and inflammation-driven remodelling of the hepatic microenvironment^1^. Characterising the molecular programmes underlying these shifts can help contextualise disease mechanisms and inform biomarker discovery.

To characterise cellular dynamics along the trajectory, we performed cell-type deconvolution of bulk liver transcriptomes (Methods; **Supplementary Fig. 8**; **Supplementary Table 14**). Hepatocytes remained the dominant cell type but progressively declined from early to late stages (from 0.66 in SW1 to 0.52 in SW13), consistent with increasing contributions from other hepatic and immune populations. To obtain robust estimates, hematopoietic cells were aggregated into myeloid and lymphoid lineages, excluding macrophages, which showed a distinct and progressive increase, particularly after mid-progression (SW6; **Fig. 4A; Supplementary Fig. 8**). Lymphoid cells (predominantly T cells and NK/NKT cells) increased more markedly than other myeloid populations, alongside rising cholangiocyte and fibroblast signals, reflecting heightened inflammatory and fibrogenic activity. In this analysis, the “fibroblast” annotation predominantly reflects hepatic stellate cells, which comprise ∼70% of this category in the reference single-cell atlas^68^. In contrast, endothelial cell proportions remained relatively stable, suggesting preservation of vascular structure even in advanced disease.

**Figure 4.**
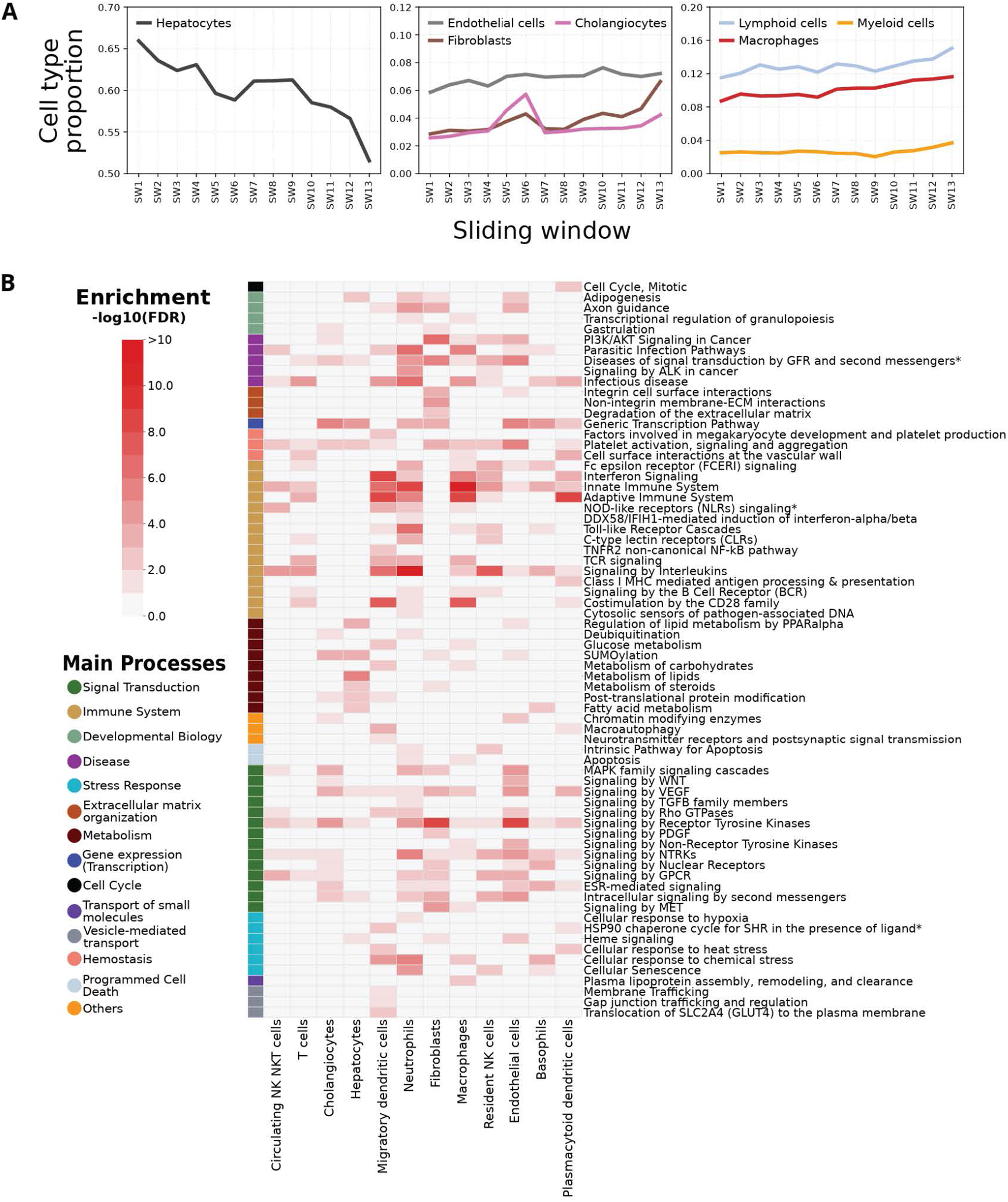
Results of cell type deconvolution analysis. **A.** Cell type deconvolution of patient transcriptomic data along our MASLD trajectory **B.** Prediction of cell types with which the deregulated processes are associated after excluding gene sets of these processes that could have been identified as differentiated by the mere change in abundance of cell types.

We next linked cell types to deregulated processes along the trajectory. To distinguish true process dysregulation from shifts in cell composition, we removed genes whose expression changes could be explained by changing cell proportions, using pseudo-bulk profiles derived from healthy liver single-cell data (Methods).

Enrichment analysis of the remaining gene sets revealed cell type–specific pathway associations (**Fig. 4B**; **Supplementary Table 15**). As expected, lipid metabolism was primarily hepatocyte-associated, immune pathways mapped mainly to macrophages, neutrophils, and migratory dendritic cells, and extracellular matrix and fibrotic processes were strongly linked to fibroblasts (**Fig. 4B**). Non-immune signalling pathways, including Wnt, receptor tyrosine kinase, and Rho signalling, were predominantly associated with endothelial cells, with additional contributions from fibroblasts, cholangiocytes, and immune cells. Together, these results highlight coordinated crosstalk between parenchymal, stromal, and immune compartments during MASLD progression.

### MASLD Trajectory-Specific Biomarkers Through Integrated Plasma-Liver Expression Analysis

To address the critical need for non-invasive diagnostic methods for MASLD/MASH staging, we sought to identify blood-based proteomics markers that predict a patient’s MASLD stage. Govaere et al.^69^ recently identified 194 genes with correlated expression in both blood plasma and liver across MASLD progression stages. We hypothesized that the 57 genes among these that were also present in our MASLD network are more likely to be directly involved in disease progression and are, thus, better suited for disease monitoring (**Fig. 5A; Supplementary Table 1**).

**Figure 5.**
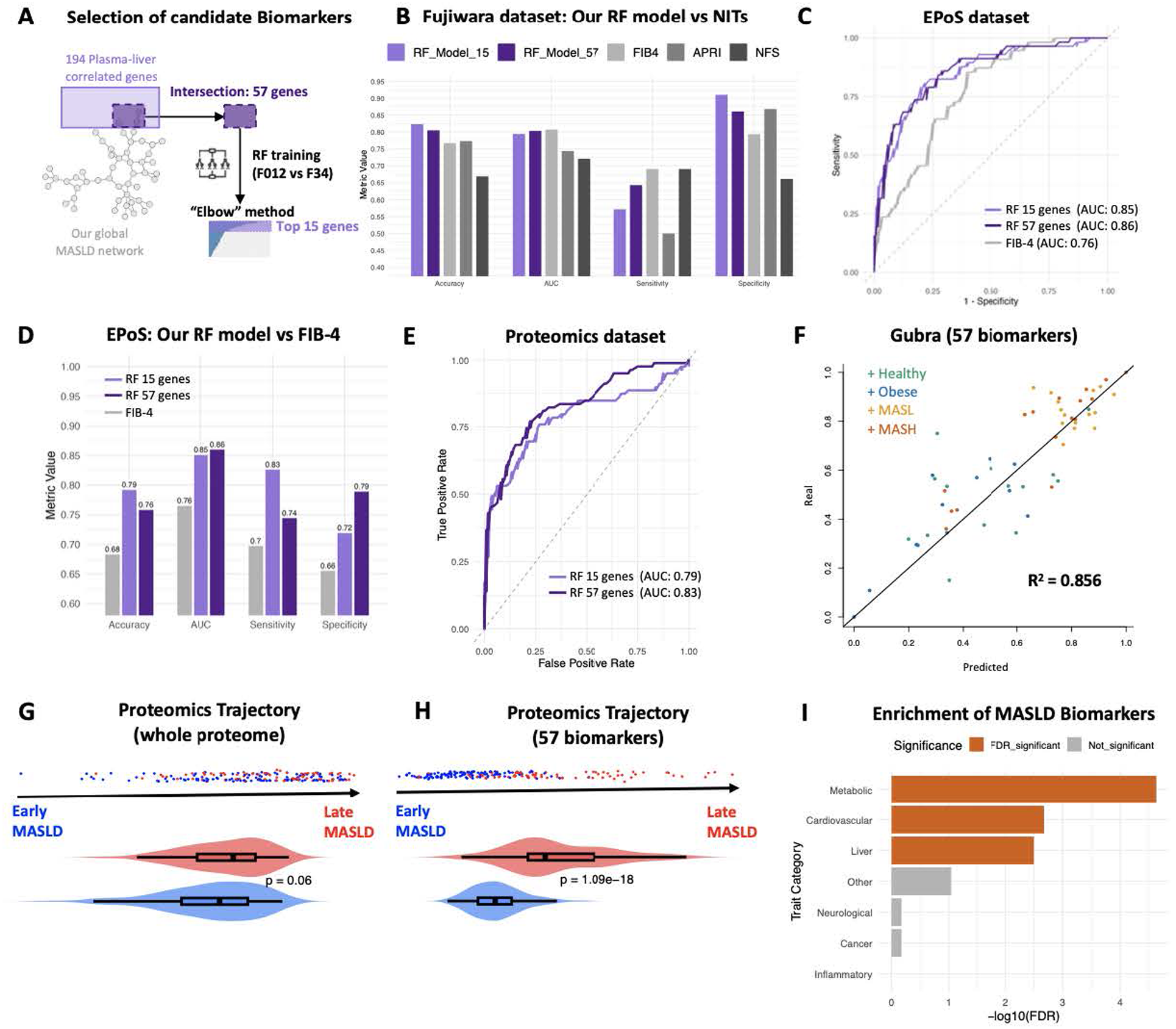
Biomarker selection and performance on the external liver transcriptomics and plasma proteomics datasets. **A.** Selection of candidate biomarkers. A 57-gene set was defined by intersecting the global MASLD network with plasma–liver correlated genes^69^, followed by random forest classification of fibrosis stage (F0–2 vs F3–4). Feature importance and the elbow method identified a reduced 15-gene subset. **Β.** Performance of the 57-and 15-gene classifiers compared with non-invasive clinical scores in the Fujiwara cohort. **C-D.** Benchmarking of the same models against FIB-4 in the EPoS cohort, with C showing the ROC curve for our model compared to the FIB-4 score, and D showing the performance using additional metrics. **E.** ROC curve for fibrosis classification of patients using external plasma proteomics data. **F.** Random forest regression predicting patient positions along the MASLD trajectory in the Gubra cohort; predicted positions (57 genes) versus transcriptome-derived positions are shown (R², Pearson correlation). **G-H.** Inferred patient trajectory positions on the external validation plasma proteomics dataset, using either the whole proteome (G) or only the 57 biomarkers (H). **I.** GWAS trait enrichment of the 57-gene biomarker panel; significant categories after FDR correction are shown in orange (Fisher’s exact test on the biomarkers associated with each of those categories, against the whole GWAS as the background).

We trained a Random Forest classifier using the 57-gene set to predict fibrosis stage (F0–F2 vs F3–F4), achieving 86.2% accuracy in the UCAM/VCU cohort (AUC = 0.769). Applying the elbow method to feature importance yielded an optimal 15-gene subset, which might be more tractable for clinical application (**Fig. 5A; Supplementary Table 1**). Validation in two independent cohorts showed robust performance. Specifically, in the Fujiwara cohort^35^, the model achieved an AUC of 0.803 (0.794 for 15 genes), outperforming established clinical scores (FIB-4, APRI, NFS; **Fig. 5B**), while in the EPoS cohort^13^ it reached an AUC of 0.86 (0.85 for 15 genes), exceeding FIB-4 by 10% (**Fig. 5C**). In general our model outperformed FIB-4 in all evaluation metrics tested (**Fig. 5D**). Robustness analysis against 300 random 57-gene signatures confirmed superior performance of the curated gene set in both external datasets (p < 1e-64; **Supplementary Fig. 9A**).

We further tested model generalisability by applying the transcriptomics-trained classifier to plasma proteomic data, where it retained strong performance (AUC = 0.83; 0.785 for the 15-gene subset; **Fig. 5E**, **Supplementary Fig. 9Β**; Methods). Together, these results demonstrate that the plasma-based gene signature generalises across cohorts, outperforms established non-invasive tests, and directly translates into blood proteomics.

To enable continuous patient positioning along the disease trajectory, we trained a random forest regression model using the 57-gene signature. In the UCAM/VCU cohort, the model achieved strong and consistent predictive performance (R² = 87.9%, p = 1.66e-08; MSE = 0.008 across 1000 iterations of 5-fold cross-validation), accurately recapitulating increasing MASLD severity (**Supplementary Fig. 9C–D**). Validation in the independent EPoS cohort^13^ confirmed this performance (R² = 85.1%, p = 8.04e-62), with predicted positions tracking progressive MASLD stages (**Supplementary Fig. 9E**), whereas random 57-gene signatures did not (R² = 0.14; Supplementary Fig. 9F).

Further validation in the Gubra dataset^34^ (Methods) demonstrated similar accuracy (R² = 85.6%, p = 1.4e-26), correctly separating healthy/obese individuals from MASLD/MASH patients along the trajectory (**Fig. 5F**, **Supplementary Fig. 9G**), with 89.2% of early-trajectory individuals classified as healthy/obese and 89.7% of late-trajectory individuals as MASLD/MASH. Finally, applying the same model to plasma proteomics data successfully recapitulated disease progression at the protein level (p = 1.09e-18; **Fig. 5G–H**). Together, these results demonstrate that the 57-gene signature robustly predicts MASLD progression across cohorts and omics layers, supporting its potential for non-invasive staging and longitudinal monitoring.

To assess genetic support for the 57-gene MASLD biomarker panel, we queried the GWAS Catalog (v1.0)^70^. Forty-seven of the 57 genes were associated with at least one complex trait, predominantly related to metabolic, cardiovascular, and liver phenotypes (**Supplementary Fig. 10A–B; Supplementary Table 16**). Nearly half of the genes (27/57; 47.4%) were linked to multiple trait categories, suggesting pleiotropy. Enrichment analysis confirmed significant overrepresentation of metabolic (FDR = 2.33e-05), cardiovascular (FDR = 0.0021), and liver-related traits (FDR = 0.0032) relative to background (**Fig. 5I**), providing genetic support for the biological relevance of the biomarker panel.

To explore potential therapeutic intersections, we mapped approved and investigational compounds to the 57-gene biomarker panel (**Supplementary Table 17**). This revealed drug–target associations spanning core MASLD processes. For example small molecules targeting COL3A1 had indications related to extracellular matrix remodelling and fibrotic disorders (e.g. Dupuytren contracture, Peyronie disease), those targeting CXCL9 and SERPINE1 are indicated for inflammatory and immune signalling-related conditions (e.g. chronic bronchitis or Alzheimer’s disease as well as various cancers). Small molecules with indications for coagulation and vascular diseases, such as atrial fibrillation, acute coronary syndrome or chronic kidney disease, target F11 and AGT, and metabolic or hepatocellular stress pathway-related indications were linked to ACAT2 and GPC3. Although exploratory, these patterns reinforce the biological relevance of the biomarker panel and illustrate how a trajectory-and network-informed framework can support hypothesis generation for biomarker-guided, stage-aware therapeutic prioritisation in MASLD. Together, these results define a network-anchored and genetically supported biomarker framework that captures MASLD progression as a continuous molecular trajectory and enables robust, non-invasive stratification of disease stage across cohorts and molecular layers.

## Discussion

MASLD has become a leading cause of chronic liver disease and liver transplantation worldwide^71^, yet its clinical assessment and mechanistic interpretation still rely primarily on static, histology-defined stages. These categories only partially capture the continuous and heterogeneous nature of disease progression and provide limited insight into the regulatory processes that govern transitions between states. Here, we present a data-driven framework that reconstructs a population-level molecular continuum of MASLD progression from cross-sectional liver transcriptomes. This trajectory is not a simulation or patient-level predictor, but a statistical representation of shared molecular states aligned with histological phenotypes across cohorts, offering a scalable alternative for studying disease progression when longitudinal biopsy data are limited or unattainable.

The inferred trajectory showed strong concordance with steatosis, ballooning, inflammation, fibrosis, and NAS and identified a robust 145-gene signature predictive of histological scores that was reproducible across multiple independent datasets, including a large multi-centre cohort and a longitudinal dataset with paired biopsies. Although MASLD progression is not strictly unidirectional at the individual level, the reproducibility of this axis across cohorts supports the existence of a dominant molecular progression pattern shared across patients. By design, our framework emphasises common disease mechanisms rather than subgroup-specific modifiers, such as sex, diabetes status, or BMI, which were corrected for during integration or did not dominate trajectory placement in this setting. This focus highlights convergent regulatory programmes that may be broadly targetable across patient populations, while providing a foundation for future stratified analyses as larger, metadata-rich cohorts become available.

Integrating co-expression analysis with transcription factor (TF) enrichment and upstream pathway information yielded a MASLD regulatory network enriched for established disease genes and validated in an independent large cohort. While module–phenotype associations remain correlative, anchoring the network to TFs and signalling pathways improves interpretability of upstream regulatory processes, as we have previously demonstrated^36^. TF-based pathway analysis refined established MASLD biology by showing that ER stress (R-HSA-380994) and KEAP1–NFE2L2 oxidative-stress signalling (R-HSA-3299685) were selectively enriched in ballooning-and fibrosis-associated TF sets, rather than in steatosis-associated TF sets, suggesting that these stress-response programmes emerge primarily during hepatocyte injury. In addition, fibrosis-specific enrichment of TP53 regulatory signalling (R-HSA-3700989) and adherens junction interactions (R-HSA-373760) highlights transcriptional and structural remodelling processes that are less emphasised in conventional MASLD pathway analyses.

By resolving progression into overlapping sliding windows rather than discrete stages, the framework captured dynamic regulatory behaviours that traditional staging obscures. Immune-related TFs and pathways, including NF-κB and STAT signalling, were sustained across the trajectory, consistent with chronic inflammatory signalling in MASLD. In contrast, profibrotic and myogenic TFs, including MEF2 family members, exhibited a transient peak during mid-progression.

Pathway-level analysis further revealed temporally ordered activation of hypoxia responses, extracellular matrix remodelling, and growth factor signalling (e.g., WNT^72^, PDGF^73^, TGF-β^74^), alongside non-monotonic metabolic changes. Persistent upregulation of glucose metabolism across the trajectory is consistent with hepatic and systemic insulin resistance. In contrast, lipid metabolism showed early suppression followed by partial reactivation, underscoring the complexity of metabolic dysfunction during disease progression.

MASLD progression is accompanied by shifts in liver cell composition driven by cellular plasticity, including stellate cell activation and hepatocyte–cholangiocyte transdifferentiation^75^. Cell-type deconvolution showed reduced hepatocyte signal and increased contributions from immune, cholangiocyte, and fibroblast cell types, consistent with known histological changes. Macrophage signals increased along the trajectory and aligned with inflammatory pathways, supporting their established role in steatohepatitis⁵⁹^,76^. Fibroblast-associated signatures, largely reflecting hepatic stellate cells within the reference atlas, were linked to extracellular matrix remodelling, whereas metabolic dysregulation was predominantly hepatocyte-driven. Although bulk deconvolution primarily supports relative trends rather than precise cell proportions, these findings are consistent with current models of MASLD and motivate future validation using single-cell^75^ and spatial^77^ approaches.

A key contribution of this study is the derivation of a 57-gene plasma-accessible biomarker panel, anchored in the MASLD regulatory network and validated across independent cohorts and omics layers. Unlike biomarker approaches based solely on predictive feature selection, this panel prioritises mechanistic relevance by linking circulating markers to disease-associated regulatory programmes and genetic evidence. The biomarker panel outperformed established non-invasive scores, including FIB-4, APRI, and NFS, in distinguishing advanced fibrosis and enabled continuous placement of patients along the molecular trajectory. Furthermore, it aligns with established fibrosis-associated signatures, overlapping with clinically validated markers such as A2M (component of the NIS4 fibrosis test^78^), FAP (associated with PRO-C3 related fibrogenesis^79,80^) and THBS2^81,82^. Importantly, our biomarker panel resolves into mechanistic axes central to MASLD progression, including extracellular matrix remodelling, immune activation, metabolic stress, and coagulation–vascular pathways^83^.

Although these biomarkers are not yet ready for clinical application, their consistent performance across transcriptomic and proteomic datasets and their relevance to known processes related to MASLD progression, supports the value of network-informed biomarker discovery in MASLD and motivates prospective validation. In this context, the observation that these biomarkers intersect with known pharmacological targets across fibrotic, inflammatory, vascular, and metabolic pathways further highlights the potential of a trajectory-informed framework to support stage-aware therapeutic hypothesis generation.

This study has limitations. The trajectory captures shared molecular progression but cannot infer causality or predict individual disease courses. Cross-sectional data limit inference on reversibility and treatment effects, and subgroup-specific modifiers such as sex, diabetes, and medication use are not explicitly modelled. Nonetheless, the resulting MASLD network provides a structured framework for contextualising molecular dysregulation, identifying stage-associated regulatory programmes, and prioritising candidate pathways for further investigation. Incorporation of longitudinal sampling, richer clinical metadata, and single-cell or spatial data will be important next steps to refine this framework and extend it toward stratified and personalised analyses.

In summary, this work provides a unified, trajectory-based molecular framework for MASLD that organises established disease mechanisms into a coherent and reproducible progression model. By integrating regulatory networks, pathway dynamics, and non-invasive biomarkers, the framework advances beyond static staging. It, thus, offers a scalable approach for studying MASLD and other progressive diseases or biological trajectories characterised by cross-sectional molecular data.

## Materials & Methods

### Datasets

For the analyses conducted in this study, 2 different publicly available human datasets have been used: the “UCAM” dataset^33^ (consisting of 58 consecutive patients recruited at the MASH Service at the Cambridge University Hospital) and the VCU dataset^14^ (consisting of 4 obese bariatric controls without MASLD and 74 patients covering the whole MASLD spectrum). The raw transcriptomic data and relevant metadata have been downloaded from Array Express (E-MTAB-9815) and NCBI’s GEO (GSE130970), respectively. Details for both datasets are given in **Supplementary Tables 1 & 2**. “Sample_5” was removed from the analysis as it was considered an outlier (Supplementary Fig. 11A-B).

For the validation of the human trajectory inference method, we used three publicly available transcriptomics datasets (GUBRA^34,84^, EPoS^13^, Fujiwara^35^) and the Govaere proteomics dataset^69^ was used for the biomarker analysis. The GUBRA dataset consists of n=26 healthy (n=14 normal-weight, n=12 overweight) individuals recruited at the Center for Diabetes Research (Copenhagen University, Denmark) and n=31 MASLD/MASH patients recruited at the Department of Hepatology and Gastroenterology (Aarhus University Hospital, Denmark). The EPoS dataset consists of 168 MASLD/MASH patients, originates from a large cohort recruited in multiple European parts, and has been previously used by our team^85^. In addition to the expression data, the EPoS dataset also provided information on the MASLD disease stage, NAS, Fibrosis stage, and Fibrosis-4 (FIB-4) scores per patient. The Fujiwara^35^ dataset contains longitudinal transcriptomic data and metadata from 58 MASLD patients across two liver biopsies spanning the full disease spectrum. As non-invasive test (NIT) scores were not directly provided in the original published dataset, the FIB-4^5^, AST-to-Platelet Ratio Index^86^ (APRI), and NAFLD Fibrosis Score^4^ (NFS) were calculated from the available metadata using standard, previously published definitions. Only samples with complete data required for each score were included in the corresponding analyses (**Supplementary Table 18**). Finally, the Govaere proteomics dataset^69^ consists of paired plasma proteomics and liver transcriptomics data from 191 patients, including samples matched to the EPoS patients, enabling cross-omics validation of the trajectory model and biomarker analysis.

### RNA-Seq Analysis and data processing

For the RNA-Seq analysis, the *FASTQC* software was used to generate quality-control (QC) reports of individual FASTQ files (http://www.bioinformatics.babraham.ac.uk/projects/fastqc, v. 0.11.9) and reads were aligned to the human GRCh38 reference genome using hisat2^87^ (v 2.1.0). Then, the genes were counted with *HTSeq*^88^ (v 0.11.1), and the *biomaRt package*^89^ was used to map “Ensembl gene IDs” to “HGNC symbols”.

We used quantile normalisation to normalise the 2 datasets and Bioconductor’s function *COMBAT*^90^ from R package *sva* to remove the batch effect observed upon integrating them (**Supplementary Fig. 11A-B**). Sex was used as a covariate. Quality control checks validate the robustness of the analysis, showing that the gene expression in each MASLD stage, separately for each dataset, remains similar after batch effect correction (**Supplementary Fig. 11C-D**). Additional stratified correlation analysis after dividing the datasets in low (bottom ⅓), medium (middle ⅓), and high (top ⅓) gene expression, also revealed high correlation scores in the gene expression before and after batch effect correction (**Supplementary Fig. 11D**). Finally, we observe a high correlation of the first principal component of the PCA plot with the main MASLD characteristics without significant association with sex (**Supplementary Fig. 11E,F**), and a similar expression of genes previously linked to MASLD before and after the correction (**Supplementary Fig. 11G**), validating the robustness of the batch effect correction.

### Pseudo-temporal ordering of patients

The merged dataset was not expected to have multiple disconnected trajectories with a particular cyclic or (necessarily) complex topology. Thus, we selected *Slingshot*^91,92^ as the most suitable tool to infer the disease trajectory. We constructed an MST (Minimum Spanning Tree) on ordered sets of clusters to capture the disease trajectory from points on the PCA plot. The inferred horizontal trajectory was consistent with the high correlation scores between PC1 and the MASLD characteristics and explained 9% of the variance of the scaled and normalised dataset (∼55% of the pre-normalised data; **Supplementary Fig. 11A, B**). It did not show any significant correlation with potential confounders, such as age (**Supplementary Fig. 11E**). This validates our design that was intended to capture the common molecular continuum of progression across the cohort, as reflected in the histological phenotypes and independent of variables such as sex or T2DM. Thus, this dimension could be considered as a pseudo-temporal trajectory of MASLD, in which low values correspond to healthy or early disease stages, as reflected in biopsy histopathology, while large values correspond to severe fibrosis conditions. For the independent dataset, this process was repeated using the complete transcriptome and only the 145 gene signatures identified by the random forest classification (see section below). The order of patients in our trajectory is described in **Supplementary Table 1.**

### Patient stratification for annotation of trajectory

Based on the histological scores defined by the CRN scoring system^6^ and following the older definition of NAFLD/NASH, our dataset (136 patients) has been divided into 4 bariatric obese controls and 132 MASLD, which has further been sub-clustered based on the fibrosis stage into mild (MASH F0), moderate (MASH F1-2) and severe (MASH F3-4) groups. This stratification aims to increase the interpretability of the analysis, which has been described in detail in our previous studies^33,85^ (**Supplementary Table 1**).

### Extraction of 145 gene signatures using Random Forest classification

By calling the parameter *rotation* from *prcomp* function in R (default stats package), we produced a list of ranked genes with the highest absolute values that explain the variance, along with the disease progression on the horizontal axis. The top 200 genes were selected as the training set of a random forest classification model (R package *randomForest*) to predict the disease stage (MASL, MASHF01, MASHF2, MASHF34) and the different MASLD characteristics (steatosis, inflammation, ballooning, fibrosis). The accuracy was estimated using 5-fold cross-validation 1000 times. From this first model run, only genes with positive “Mean Decrease Accuracy” were selected, yielding the set of 145 top genes that contributed to disease stage prediction (**Supplementary Table 1**). Those genes were used as the training set of a new random forest classification model. Testing the model on three independent cohorts (GUBRA^34,84^, EPoS^13^, Fujiwara^35^) conclusively demonstrated its generalisability. **Supplementary Table 3** shows AUC values from ROC plots for two-way classification of the groups, as we cannot generate 3-way classification ROC.

### Pathway enrichment analysis

The Reactome Pathway Knowledgebase^38^ (accessed May 2024) was used in different steps of this study to perform pathway enrichment analysis. Fisher’s exact test^93^ was used for the statistical significance test, and the Benjamini–Hochberg method^94^ was applied for the multiple hypothesis testing correction. The KEGG database was used for pathway enrichment analysis of the 145 gene signature in Figure 1E to facilitate visualisation, as these pathways are more generic.

### Identification of gene expression modules associated with MASLD histological phenotypes

We performed weighted gene co-expression network analysis (WGCNA)^95^, as in Barker et al.^36^, using the R package *WGCNA*^37^ (v 1.72-1) on the full transcriptome to capture MASLD-related molecular programmes. A signed network was constructed using a soft-thresholding power of 4, the minimum value that achieved an approximate scale-free topology (fit index = 0.9). Adjacency weights were transformed into a topological overlap matrix (TOM), and gene co-expression modules were identified from the resulting dissimilarity matrix (1–TOM) using dynamic tree cutting (function *cutreeDynamic,* package *dynamicTreeCut* v1.63-1).

Module–phenotype associations were assessed by linking module eigengenes to histological traits, with inflammation included as a covariate to control for its dominant transcriptomic signal and its ambiguous role as cause versus consequence of disease. As inflammation is incorporated into the NAS score, this approach enables identification of modules specifically associated with steatosis, ballooning, fibrosis, and NAS beyond global inflammatory effects. We also included age as a covariate because it is a well-established determinant of MASLD progression, influencing both hepatic and systemic metabolic states.

To optimise module detection, we tuned the *deepSplit* and *minClusterSize* parameters using a cross-validated optimisation strategy. Module eigengenes were evaluated for their ability to model the four MASLD phenotypes using linear regression (*lm* and *step* functions of the *stats* R package) with four-fold cross-validation (*crossv_kfold* function of the *modelr* R package (v 0.1.11)), selecting parameters that maximised predictive performance while adjusting for inflammation.Each linear model had the following format:

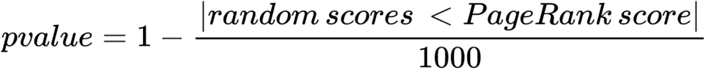

We optimised linear models using backward stepwise selection with the Akaike Information Criterion (AIC)^96^, retaining only modules with significant coefficients (adjusted p < 0.05, Benjamini–Hochberg correction^94^). Model performance was evaluated on held-out test sets using mean squared error (MSE). Average test MSEs across the four MASLD phenotypes were used to identify optimal parameters, resulting in the following parameter settings: *deepSplit* = 4 and *minClusterSize* = 180. Final models were built using these parameters. For the downstream analysis, we filtered the module gene sets based on their Pearson correlation with the corresponding eigengene. For modules with fewer than 1,000 genes, we removed genes with non-positive correlation. For larger modules, we applied a cutoff at the 20th percentile of the correlation scores to exclude genes with lower values (**Supplementary Table 6**).

### Transcription factor activities analysis

We used *VIPER*^97^ (Virtual Inference of Protein-activity by Enriched Regulon analysis; v 1.37.0) and the CollecTRI database^98^ (accessed May 2024) to identify the upstream transcriptional regulators that can explain the observed gene expression changes. The calculated Normalised Enrichment Score (NES) was used to determine the direction of each regulation (“activation” for TFs with NES > 0 and “inhibition” for those with NES < 0). We performed hypothesis testing using the Student’s t-test, and then, the Benjamini-Hochberg method^94^ was used to correct the p-values, thereby reducing Type I errors (**Supplementary Tables 8 & 12**).

### Constructions of global MASLD network

To construct a global interaction network capturing molecular processes across MASLD stages, we integrated signalling and transcriptional regulation inferred from WGCNA modules, differential expression, and TF activity across SWs. For each MASLD phenotype, we generated upstream (signalling) and downstream (transcriptional) subnetworks using the *Prize-Collecting Steiner Forest* (PCSF) algorithm^99^ (*PCSF* R package v0.99.1), and subsequently merged them into a unified MASLD network.

*PCSF* returns as output a subset of a given reference network relevant to an input-weighted list of prize-carrying nodes. Prize-carrying nodes were defined from two complementary sources. First, for WGCNA-derived modules, we performed TF enrichment using CollecTRI^98^ and TF-regulon databases from *EnrichR*^100^ (ENCODE and ChEA Consensus TFs from ChIP-X^101^, DNA binding preferences from JASPAR^102,103^, TF protein-protein interactions), followed by Reactome^104^ pathway enrichment of the enriched TFs to identify semantically specific signalling pathways (**Supplementary Tables 8 and 9**). Filtered module genes (downstream nodes), enriched TFs, and genes from associated Reactome pathways (upstream nodes) constituted the prize-carrying nodes for each MASLD variable.

Second, for differentially expressed genes, we selected TFs deregulated in at least one SW (FDR < 0.05; **Supplementary Table 12**), identified their associated signalling pathways (**Supplementary Table 9,** as above), and intersected their regulons (using the CollecTRI annotation^98^) with WGCNA modules. These regulon genes, deregulated TFs, and pathway genes defined the prize-carrying nodes for the deregulated network. All enrichment analyses used Fisher’s exact test with Benjamini–Hochberg correction (FDR < 0.05), restricting background genes to those detected in the transcriptomic data.

The reference interaction network was assembled by integrating curated signalling and metabolic interactions from Reactome^104^, KEGG^105,106^, HumanCyc^107^, ReconX^108^, provided by Pathway Commons^109^ (v12), and OmniPath^110^, accessed using the *OmniPathR* library (v3.17) in Bioconductor^111,112^ and using the *union* function of the *igraph* R package (v 2.03)^113^. Weakly supported OmniPath interactions were removed, keeping only those where the number of resources and curation efforts were above the 0.5 and 0.75 quantiles of their respective distributions. Dead-end metabolites were also excluded, protein complexes were expanded into fully connected subgraphs, and inhibitory edges were discarded. The resulting reference network contained 12,549 nodes and 205,706 edges (**Supplementary Table 19**).

PCSF was run using fixed edge costs (0.001) and node prizes (1). For each of the five MASLD variables, upstream and downstream subnetworks were generated and iteratively connected, using a randomized PCSF variant (10 iterations adding random noise) to improve robustness. This gradual generation of the MASLD network was selected to preserve information about the origin of each node and edge. These subnetworks were then merged into a global MASLD network, using the *union* function of the *igraph* R package (v 2.0.3).

To refine the network, we retained only functionally annotated genes and weighted edges based on Gene Ontology Biological Process^114^ semantic similarity, computed using the *GOSemSim* R package^115^ (v 2.26.0) and averaged across Resnik^116^, Lin^117^ and Wang^118^ metrics. Nodes lacking GO annotation were removed, and Laplacian normalisation was applied to reduce hub bias^119^. The final MASLD network comprised 7,165 nodes and 69,314 edges (**Supplementary Table 10**).

Network relevance was assessed by enrichment against two independent MASLD gene signatures: a literature-curated set (164 genes^83,120–124^; **Supplementary Table 11**) and the WikiPathways MASLD pathway from MSigDB^55^ (155 genes; 15-gene overlap). Enrichment significance was evaluated using Fisher’s exact test.

### Patients’ stratification into sliding windows

We divided the ordered samples along the trajectory into sliding windows (SWs), namely a sequence of overlapping groups [SW_1_, SW_2_,…, SW_X_], to investigate how the MASLD molecular profile changes in the pseudo-temporal space. To define an optimal SW configuration, we developed a graph-based optimisation framework that evaluates all possible sequences of SWs given a set of window sizes *S=[S_1_,…, S_m_]* and a predefined overlap percentage *α* between adjacent windows.

In this framework, disease stages are represented as nodes in a directed acyclic graph (DAG), where each node corresponds to a group of samples aligned by pseudo-temporal order. Directed edges connect overlapping windows, ensuring continuity along the trajectory. The objective is to identify a path through this graph that best captures progressive molecular changes by enabling sequential comparisons between adjacent windows. Our method is implemented using a three-step workflow: creating of all potential directed paths of SWs, evaluating and filtering, and finally, selecting the optimal one.

The method begins by ordering the N samples according to their scores on the first principal component (PC1) of the expression data. Based on this ordering, each sample is assigned a unique index i ∈ [1, 2,…, N]. Using these indices, an iterative process is performed to construct a directed acyclic graph (DAG), in which nodes correspond to groups of samples and directed edges represent a valid transition between two overlapping groups. An empty starting node is first created as the origin of all paths in the DAG (**Supplementary Fig. 3A**). In the first iteration, a set of nodes is generated using a predefined collection of window sizes *S*∈ *[S_1_,…, S_m_].* For each window size *S_i_* a node is created that contains the first *Si* samples (indices 1 through *Si*). These nodes constitute the candidate versions of SW_1_. Directed edges from the starting node to the SW_1_ nodes are created. In subsequent iterations, each existing SW node is extended forward along the index order to generate candidate nodes for the next SW. For a SW node with *K* samples, the starting index of its subsequent SW nodes is defined as *K+1-ceiling(K*α)*, where *α* controls the degree of overlap between consecutive windows and ensures that the last 100*α%* percent of the samples in the current SW will also be included in the next window. New directed edges connect the current node with its subsequent ones. This iterative growth of the graph terminates when all the samples have been mapped to SW nodes. Many nodes and edges could be generated multiple times through the expansion of different paths. Thus, filtering is performed at the end of each iteration to remove this redundancy. Finally, an empty ending node is created and connected with the final SW nodes. All directed paths connecting the starting and ending nodes constitute potential representations of the pseudo-temporal sequence of disease stages (**Supplementary Fig. 3A**).

To evaluate candidate paths, we performed differential expression analysis between each pair of adjacent SWs using DESeq2^125^. Edges were initially retained if the number of differentially expressed genes (DEGs; FDR < 0.1) fell within an empirically defined range *R=[100, 1000]*, reflecting sufficiently strong but not excessive differentiation (that would be best captured in more than two SWs). Graph evaluation proceeded iteratively. To reduce computational burden, paths containing edges outside this range were pruned at each iteration. To preserve graph connectivity, DEG thresholds (i.e. the range of *R*) were adaptively relaxed such that the lower bound did not exceed the 80th percentile and the upper bound did not fall below the 20th percentile of the observed DEG distribution.

This procedure typically yielded multiple candidate paths. Additional filtering was applied to select the optimal one. First, incoming edges to each SW node were clustered by the upstream node’s starting index, retaining only the edge with the highest DEG count (or, if tied, the one spanning more samples). Second, edges below the lower DEG threshold were used to compute a median DEG value, and edges with fewer DEGs were removed while preserving graph connectivity (**Supplementary Fig. 3B**). Remaining paths were ranked using the product of three max-scaled criteria: (i) median DEG count, (ii) number of edges exceeding the lower DEG threshold, and (iii) coefficient of variation of DEG counts along the path, using the following pairs of SW sizes and overlap scores: ([6,15], 0.2), ([8,17], 0.25), ([8,24], 0.3), ([10,20], 0.3), ([12,25], 0.35). Therefore, five graphs were constructed and filtered, and the candidate paths from all of them were prioritized to find the optimal SW sequence (**Supplementary Fig. 3C**; **Supplementary Table 5**).

## Statistical analysis

Differential gene expression analysis was done using *DESeq2*^125^ (v 1.26.0), and the Benjamini-Hochberg method was applied to adjust the raw p-values to control the False Discovery Rate (FDR). In analyses using discrete stratification of patients’ disease stages, comparisons were made to the immediately preceding stage. For all comparisons related to the trajectory analysis (see section “Patients stratification into sliding windows”), sliding windows (SWs) were compared to the previous SW (SW_x_ vs SW_x-1_).

### Disease stage-specific network analysis

We applied a network propagation-based approach to detect the deregulated components of the MASLD network for each SW using the *igraph* R package (v 2.0.3). We used the personalised *PageRank* algorithm^126^ to run network propagation. This method accepts a vector of weights for a set of nodes as input. We defined these weights based on results from differential expression analysis (for all genes except transcription factors) and TF activity analysis (for transcription factors). We were interested in finding a network signature for up-and down-regulated conditions separately, so we created two weight vectors for each SW. In particular, we defined the weights of node i (𝑊_+_ and 𝑊_-_) using the direction of its differentiation (based on the logarithmic fold change or NES score) and the log-transformed adjusted p-value of the respective statistical test, to be bounded by a maximum value of 10:

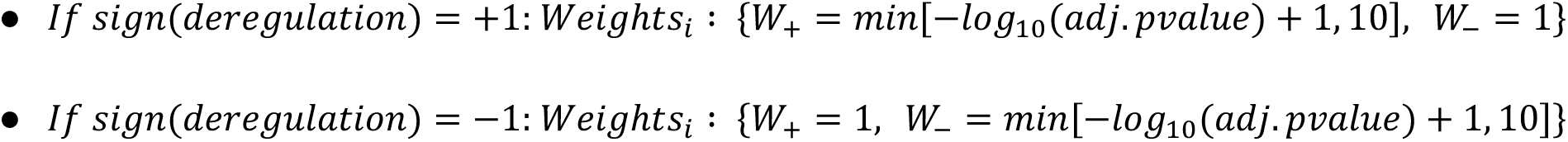

Network propagation was performed for each SW and direction of deregulation. To assess the statistical significance of the derived PageRank scores, we run the propagation task 1000 times using random weight distributions. Then, we calculated an empirical p-value for each node based on the ratio of its random scores that are lower than the real one:

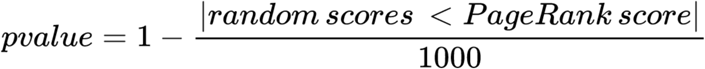

We implemented this approach for three values of the damping factor: [0.5, 0.7, 0.85], and we selected the gene nodes with p-value<0.05 in all three runs as significantly propagated. Then, we extracted the respective subnetwork signature as a subgraph of the MASLD network and we removed the isolated nodes, keeping only the components with two or more members. Finally, the activation score of a gene in a SW was defined as the average of the log-transformed empirical p-values obtained from the network propagation runs using the three damping factors.

### Definition of disease-relevant pathways

We used pathway enrichment analysis to link deregulated subnetworks in each SW to molecular pathways using Reactome^38^ (see “Pathway enrichment analysis”; **Supplementary Table 13**). This analysis yields distinct sets of significantly enriched pathways (terms) for each SW and each direction of deregulation. To extract a unified, comprehensive set of MASLD-relevant pathways with minimal semantic redundancy, we created a slim version of Reactome by pruning its hierarchical structure. We finally project all the enriched terms on that slim graph.

First, we constructed the Reactome graph using the *igraph* Python package (v 0.4.11), to encode parent–child semantic relationships. We calculated the information content^116^ (IC) and semantic value^127^ (SV) of Reactome terms using their reference genomic annotations. These two properties capture the specificity of terms within the graph, with higher values corresponding to more semantically specific terms. Firstly, we retained the most generic terms by filtering out those above the 25th percentile of each distribution. This filtering step reduced the number of Reactome terms from 2,655 to 315. Secondly, we examined the semantic distances between the leaf terms of the pruned graph and their parents terms, by calculating the differences of their IC and SV values. If both distances were lower than the median values of respective distributions and the parent term was not a direct child of the graph root, then the child terms were filtered out, as they were considered semantically similar to their parent. This step defined the final slim version of Reactome, which consisted of 225 terms (**Supplementary Table 20**).

To define the list of MASLD-related pathways, we used the Reactome slim graph as a framework. We first performed Reactome enrichment analysis on the global MASLD network (FDR < 0.05) and intersected the enriched terms with those in the graph. This resulting subgraph was used as a reference to filter the SW-level enriched pathways (FDR < 0.05). For each set of enriched pathways, we checked whether they were present in the reference subgraph or could be mapped to ancestral terms within it. Enriched terms that failed to be mapped to the reference subgraph were removed. Additionally, terms with fewer than 10 genes in the respective SW-based deregulated subnetworks were removed as they were considered highly specific for the comprehensive set of MASLD-related pathways. This filtering step created a MASLD-related subgraph of Reactome with 141 pathways (**Supplementary Table 20**). Finally, we selected the leaf terms of that subgraph, along with their parental terms if they were not directly linked to the root and the annotations of their child terms did not capture more than 20% of their genomic annotations in the deregulated networks. This final step defined a list of 111 MASLD-related pathways (**Supplementary Fig. 6**; **Supplementary Table 13**).

To calculate the activation scores of the MASLD-related pathways along the disease trajectory, we used the activation scores of their related genes sets ((see “Disease stage-specific network analysis”). We created a subnetwork for each pathway, intersecting the global MASLD network and their reference gene sets. The activation scores of each pathway for a specific SW was defined as the summation of the activation scores of its related genes in that SW, normalised by their pagerank centralities^126^ on the pathway-based subnetwork. The respective cumulative scores were defined by progressively summing the SW-based activation scores (**Supplementary Fig. 6**; **Supplementary Table 13**).

### Cell-type deconvolution analysis

As a reference for the deconvolution analysis, we used the Liver Cell Atlas, a 10X single-cell RNA-seq dataset from a healthy human liver^68^. We re-analysed the single-cell dataset following standard single-cell analysis workflows^128^. Genes expressed in fewer than three cells and cells expressing fewer than 200 genes were removed. Cells with more than 7,000 detected features (putative doublets) or with >5% mitochondrial gene content were also excluded. The filtered data were then normalised and variance-stabilised, followed by identification of highly variable genes (HVGs), principal component analysis (PCA), and UMAP dimensionality reduction to assess data structure and suitability for downstream analyses.

For the deconvolution analysis, we applied the CATD pipeline^129^, using the raw counts as input. CATD performs normalisation and additional preprocessing internally. After setting the preprocessing parameters, we run the deconvolution analysis, selecting a subset of methods (EpiDISH^130^, bseqsc^131^, DWLS^132^, CIBERSORT^133^, FARDEEP^134^ and BayesPrism^135^). The consensus result was calculated as the mean of cell-type proportion per cell-type across methods (**Supplementary Table 14**). We used these methods to calculate the consensus cell type proportions because they ranked among the top-10 of the 31 deconvolution methods of the CATD pipeline across multiple metrics, based on benchmarking on real bulk RNA-seq samples^129^.

### Identification of cell-type specific deregulated pathways

We assumed that differentiated network signatures arise from either shifts in cell-type composition or from deregulation of cell-intrinsic biological processes. To disentangle these effects and identify pathway deregulation attributable to specific cell types, we applied the following workflow:

Initially, using patient stratification into SWs and the corresponding cell-type proportions (see “Cell type deconvolution analysis”), we modeled Gaussian kernel density functions for cell-type proportions within each SW. These kernels were used to generate pseudo-bulk RNA-seq samples by sampling cells from a healthy liver single-cell reference dataset^68^, with predefined library sizes. For each SW, the number of pseudo-samples (6–20) and library sizes (20–50 million counts) were randomly selected. Differential expression, TF activity, and network-based analyses were then performed on these pseudo-datasets using the same pipeline as for the real data, under the assumption that the resulting signatures reflect network differentiation driven solely by changes in cell-type composition. We generated 30 pseudo-datasets and averaged differential expression and TF activity results across replicates.

Network analysis of the pseudo-datasets yielded up-and down-regulated components, which were compared with the corresponding real SW signatures. Genes present in both real and pseudo signatures, whose activation score in the pseudo-signature exceeded that in the real signature, were classified as composition-driven and removed from MASLD-relevant pathways (see “Definition of disease-relevant pathways”). The remaining genes were therefore assumed to reflect pathway deregulation independent of changes in cell-type proportions.

To assess whether deregulated pathways were associated with specific liver cell types, we first derived extended cell-type marker sets from the reference single-cell dataset using the *Seurat* R package (v. 5.0.3)^136^. Marker lists were filtered to retain only genes present in the global MASLD network. Each marker set was then expanded using the PCSF algorithm (with the same parameters as in “Generation of global MASLD network”; *PCSF* R package v0.99.1) to incorporate topologically proximal genes from the MASLD network and amplify each gene set. Finally, enrichment analysis was performed to evaluate the association between deregulated pathways and cell types using Fisher’s exact test, with Benjamini–Hochberg correction for multiple testing (FDR<0.05; **Supplementary Table 15**).

### Machine Learning Models for Non-Invasive MASLD Biomarker Prediction

#### Random forest classification for fibrosis stage prediction

Govaere et al, identified a set of 194 genes whose gene expression in the liver correlated with that at the proteome plasma level^69^. In brief the data for this study came from 191 histologically characterized NAFLD cases collected across six European specialist centres participating in the European NAFLD Registry (NCT04442334), encompassing fibrosis stages F0–F4 and centrally scored using the NASH-CRN system. Protein quantification was performed using the SomaScan aptamer-based platform (SomaLogic).

From this set, we derived a curated panel of 57 biomarker genes by intersecting the 194 genes with the differentiated component of our global MASLD network. A random forest classifier was trained to distinguish early (F0-F2) from advanced fibrosis (F3-F4). Training was performed on the UCAM/VCU internal cohort using 5-fold cross validation across 1000 iterations to ensure model stability. Gene-importance scores were extracted, and the “elbow” method was applied to select a subset of 15 genes above the inflection point. A second classifier was trained using this gene subset.

Performance was assessed using ROC analysis, area under the curve (AUC), sensitivity, specificity, and overall accuracy as evaluation metrics. External validation was performed on the EPoS^13^ and Fujiwara^35^ datasets, with the classifier trained exclusively on the UCAM/VCU dataset and applied to those two external datasets without retraining. For benchmarking, we directly compared its performance with commonly used non-invasive fibrosis scores used in clinical practice, including FIB-4, APRI, and NFS (when available). To assess model robustness we generated 300 random 57-gene sets and compared their AUC distributions with that of our curated 57-gene panel.

To test translatability from transcriptomics to proteomics, we applied the trained model to the plasma proteomics measurements from Govaere et al ^69^. The same 57-gene (protein) signature was used, and corresponding protein abundance was z-score-scaled prior to prediction. Model predictions were compared to patient fibrosis stages (F0–F2 vs F3–F4) using ROC analysis. The resulting AUC, sensitivity, and specificity were calculated at the optimal threshold determined by Youden’s J index. We acknowledge that the plasma proteomics data were taken from Govaere et al.^69^, who identified liver–plasma correlated proteins; however, no feature selection or model training was performed on these data. All model development was carried out exclusively in independent liver transcriptomic cohorts, with proteomics used only to assess cross-omic translatability rather than independent discovery or validation.

#### Random forest regression for trajectory position prediction

To predict each patient’s position along the MASLD trajectory, we developed Random Forest regression models trained using our 57-and 15-gene signatures, respectively. For each model, the data were randomly split into training (80%) and testing (20%) sets, and performance was evaluated across 1000 iterations by calculating the mean R^2^ and its corresponding p-value (Pearson correlation between full-transcriptome trajectory positions and predictions based on the 57 or 15 gene sets).

Expression data were transformed into z-scores based on the 194 gene set (194 plasma proteins correlated with liver transcriptomics according to the proteomics study from Govaere et al^69^). This approach ensures predictions even when partial proteomic measurements are available, enabling potential real-time prediction of patients’ position on our trajectory based solely on the most informative plasma proteins.

The same regression models were applied on two independent transcriptomic datasets (Gubra and EPoS) and to the Govaere plasma proteomics dataset^69^, using the UCAM/VCU model for training and the external datasets exclusively for testing. Generalisability was confirmed by comparing predicted vs reference trajectory positions. Random 57-gene sets were used as negative controls, evaluated over 300 runs per dataset, with predictive metrics averaged, highlighting the predictive value of our selected gene signature.

### GWAS Catalog mining and trait categorisation

The GWAS Catalog (v1.0)^70^ was used to extract all associations between our selected biomarkers and traits (**Supplementary Table 16**). For each gene, all associated GWAS traits were aggregated into biologically interpretable categories using regular expressions. The predefined categories were: Metabolic, Liver, Cardiovascular, Inflammatory, Neurological, and Cancer, while traits that did not match any of these were assigned to “Other”. The number of unique genes per category was calculated and visualised using bar plots. Genes mapped to multiple categories were identified and reported separately. Gene–trait relationships were visualised using a dot plot, illustrating the breadth of cross-system associations. To test whether specific trait categories were overrepresented among the 57 MASLD biomarkers relative to the background set of all genes reported in the GWAS Catalog, Fisher’s tests were performed per category. Finally, p-values were adjusted using the Benjamini–Hochberg method (FDR < 0.05).

### Mapping of small molecules onto biomarker panel

To identify potential candidate small molecules that can be used to target our biomarkers we searched in the ChEMBL database^137^ for approved or clinical candidate drugs (max_phase: [2,3,4]) that target these genes. Each biomarker had been identified as up-or down-regulated in specific SWs based on our network analysis. We therefore searched for drugs whose action potentially inverses the deregulation of the respective targets (agonists/activators for the down-regulated and inhibitors/antagonists for the up-regulated genes). Finally, we created a mapping of drugs and drug targets for molecular events related to the 10 biomarkers for which small molecules targeting them existed in the database (**Supplementary Table 17**).

## Data and code availability

All data used in this study are publicly available as described in the methods. The code used for the analyses can be found here: https://github.com/kamzolas/MASLD---Continuous-trajectory-approach

## Acknowledgements

The work was supported by the European Molecular Biology Laboratory (IK, TK, CGB, AVP, IP, HW, EP) and Open Targets (TK, IP, EP); The EMBL GeneCore Facility is acknowledged for support in transcriptomics data collection. EMBL IT Support is acknowledged for the provision of computer and data storage servers. Funding from the Medical Research Council (MRC) and the NIHR Cambridge Biomedical Research Centre (NIHR203312) supported I. Kamzolas. MRC MDU, Caixa Foundation and Spanish Government ATRAE programme support AVP research on MASLD.

## Author contributions

IK, TK: Conceptualization, Methodology, Software, Validation, Formal analysis, Investigation, Data curation, Writing - Original Draft, Writing - Review & Editing, Visualisation. CGB, AVP: Formal analysis, Investigation, HW: Data Curation, MV: Writing - Review & Editing, Supervision, NF, YH, QMA: Resources, IP: Supervision, Funding acquisition, AVP: Conceptualization, Supervision, Resources, Funding acquisition, Writing - Review & Editing EP: Conceptualization, Methodology, Resources, Supervision, Project administration, Visualisation, Funding acquisition, Writing - Original Draft, Writing - Review & Editing.

## Supplementary Figure legends

**Supplementary Figure 1. MASLD patients along the continuous transcriptomics-driven trajectory for the internal UCAM/VCU dataset. A.** The colouring is based on the histopathologists’ scores. **B.** Triangles showing the average horizontal position of each scoring group on the trajectory. Both panels provide information for the NAS score (ranging from 0 to 7), Steatosis scoring (0-3), Inflammatory damage (0-3), Ballooning (0-2), and Fibrosis (0-4). **C**. Violin plots illustrate the distribution of the patients in each histopathological scoring group. Each panel shows the Pearson correlation coefficient (R score) for the mean trajectory positions of each scoring group. The p-values are the results of ANOVA followed by Tukey’s pairwise comparisons for the distributions of patients across the histopathological scores. The statistically significant groups are indicated with dashed lines (*p-value <.05, ** p-value <.005, *** p-value <.0005).

**Supplementary Figure 2. Trajectory inference on independent MASLD transcriptomic datasets and MASLD patients along the continuous transcriptomics-driven trajectory for the external longitudinal (Fujiwara) dataset. A.** Trajectory inference for the independent EPoS dataset using the whole transcriptome. Color coding depicts MASLD stages as published with the original RNA-Seq data. Triangles show the average horizontal position of each scoring group on the trajectory. **B.** Trajectory inference for the independent GUBRA dataset. The population was characterised as Healthy (non-obese individuals without MASLD/MASH), Obese (healthy but obese individuals without MASLD/MASH), MASL, and MASH. **C.** Fujiwara dataset: Patient movement on the trajectory between the two biopsies. Movement towards the left or right indicates disease regression or progression, respectively. The colour shows the change in NAS score between the two biopsies, with positive scores indicating an increase in the second biopsy. **D.** Overview of patients’ positions shift on the trajectory by the NAS change between the two biopsies. Statistical significance between the patients’ position change for the increased and decreased NAS score was calculated using the Wilcoxon test. **E.** Similar to (D) but for the change in fibrosis score.

**Supplementary Figure 3. Graph-based approach to find the optimal sequence of sliding windows for the stratification of patients. A.** Generation of a directed acyclic graph, where each node corresponds to a group of samples (sliding window; SW) aligned by pseudo-temporal order. Directed edges connect overlapping windows, ensuring continuity along the trajectory. The graph is generated iteratively until all the samples have been mapped to SW nodes. **B.** Pruning of the graph to identify candidate SW sequences: differential expression analysis is performed for each linked pair of SWs on the graph, retaining only edges that meet specific criteria regarding the number of differentially expressed genes (FDR < 0.1). **C.** The optimal SW sequence is defined as the one that maximizes the product of three max-scaled criteria: (i) median DEG count, (ii) number of edges exceeding the lower DEG threshold, and (iii) coefficient of variation of DEG counts along the path.

**Supplementary Figure 4. Agglomerative clustering of the 57 TFs that were found to be deregulated in two types of discrete patient stratification (Mild, Moderate, Severe and NAS scores), and the pseudotemporal trajectory**. Reactome pathway enrichment analysis was performed to link TFs to molecular pathways and to create the corresponding one-hot encoding matrix. The clustering was performed using this matrix to identify groups of functionally related TFs.

**Supplementary Figure 5. Correlation plots of the activities of TFs for the UCAM/VCU and EPoS datasets**. **A.** Pearson correlation of cumulative activation scores of TFs, which were identified as deregulated (FDR<0.05) based on the sliding windows analysis in at least one dataset. **B.** Pearson correlation of the cumulative TF activation scores between the two datasets for each sliding window. Each dot represents a TF identified as deregulated (FDR<0.05) at least in one sliding window of a dataset.

**Supplementary Figure 6. Cumulative activation scores of Reactome pathways along the pseudotemporal trajectory.** This list of terms provides a comprehensive description of MASLD-relevant pathways, derived from the enrichment analysis of deregulated network signatures along the disease trajectory and a pruning procedure applied to the hierarchical structure of Reactome. The SW-based activation score of each pathway was defined as the sum of the activation scores of its related genes, normalised by the pagerank centralities of these genes within the pathway-based subnetwork. The respective cumulative scores were defined by progressively summing the SW-based activation scores.

**Supplementary Figure 7. Correlation plots of the activities of pathways for the UCAM/VCU and EPoS datasets**. **A.** Pearson correlation of cumulative activation scores of Reactome pathways that were identified as MASLD-relevant, based on the sliding windows analysis in at least one dataset. **B.** Pearson correlation of cumulative activation scores of Reactome pathways between the two datasets for each sliding window. Each dot represents a pathway identified as MASLD-relevant at least in one dataset.

**Supplementary Figure 8. Consensus proportion of cell type along the disease trajectory**. Cell type deconvolution was applied on the bulk RNA-seq samples using 6 methods. The average cell type proportions for each sliding window were calculated by averaging the proportions across the corresponding samples. Finally, the consensus result was calculated as the mean of cell-type proportion per cell-type across methods.

**Supplementary Figure 9: Performance of the random forest classification and regression models. A.** Performance of model using the 57-biomarker set compared to using random genes. In both external validation datasets (EPoS and Fujiwara), similar models were run 300 times, using random 57-gene signatures as input. The red points show our model’s AUC for the 57-biomarker set, while the gray points show the model’s performance for the random sets. **B.** Selection of optimal threshold using Youden’s index on the proteomics dataset. **C.** Patient distributions on the predicted trajectory using the random forest regression model for the internal UCAM/VCU dataset. Run 1000 times using 5-fold cross validation and the distribution is generated using the mean result for each patient. Different colors denote different MASLD stages. Asterisks represent statistical significance after applying ANOVA among the disease stages; *: p < 0.05, **: p < 0.005, ***: p < 0.0005, ****: p < 0.00005. **D.** The y-axis shows the positions of the patients on the “real” trajectory, inferred using the whole transcriptome. The x-axis shows the predicted positions of patients using only the 57 biomarkers. R^2^ denotes the Pearson correlation score for real vs predicted positions on the trajectory. **E.** Similar to (D) but for the external EPoS dataset. **F.** Similar to (E) but for random 57-gene signatures. **G.** Results of the random forest regression model that sorts patients on the MASLD trajectory and performance on the Gubra dataset (additional validation dataset). Mean squared error for the predicted patients’ positions on the trajectory. The different colours depict the various disease stages (Healthy, Obese, MASL, MASH). The patients are sorted by their positions along the trajectory, independent of their relative distances.

**Supplementary Figure 10: Genetic evidence of our 57-biomarker set. A.** Gene-trait category associations based on the GWAS catalog. Of those, 47/57 biomarkers were associated with at least one trait (**Supplementary Table 16**), while 13 biomarkers were part of our top 15 biomarker list (selected using the elbow method). **B.** Number of biomarkers associated with metabolic, cardiovascular, liver, cancer, neurological or inflammatory traits. Those associated with a different trait are classified as “other”.

**Supplementary Figure 11. Quality control and validation for dataset integration**. **A.** PCA plot of the merged UCAM/VCU quantile normalised counts. There is an apparent batch effect separating the patients into two clusters stemming from the dataset from which they are derived. **B.** COMBAT batch effect correction solved the problem of the batch effect with patients altogether in a cluster and a clear disease progression pattern from the right towards the later disease stages on the left. The outlier (sample 5) was removed from all downstream analyses. **C-D.** Normalised counts before and after batch effect correction, separately for each disease stage. The data suggest that COMBAT has not significantly altered gene expression, as indicated by the high correlation (R > 0.97) across all MASLD stages in the VCU (C) and UCAM (D) datasets. Inset in panel D shows the mean correlation in expression before and after correction stratified by low, medium, and high expression levels. The categories were defined by ranking genes by normalised expression in each stage and dividing them into three groups (High: top third, Medium: middle third, and Low: bottom third). The dots inside each bar correspond to the correlations of expression separately for each disease stage (dot colours in agreement with disease stage colours from panel C). **E.** Correlation of the principal components (PCs) with the variables of interest. The first 10 PCs have been correlated using Pearson correlation with 6 continuous variables of interest: age, NAS score, Fibrosis, Steatosis, Ballooning, and Inflammation. **F.** Heatmap of the quantile normalised counts before and after the merging/batch effect correction for UCAM and VCU datasets. The rows show genes previously linked to MASLD^138–141^. The data are shown in a “fold change vs the first disease stage” of each dataset, respectively (“red” = increase; “blue” = decrease; “white” = unchanged; “green datasets”: UCAM and VCU before correction; “blue datasets”: UCAM and VCU after correction; the asterisks “*” in the column names indicate that the samples have been batch-effect corrected). Note that the control group (the first column in each block) in the VCU dataset is labelled as CTRL, while in the UCAM it is labelled as MASL.

## Supplementary Table legends

**Supplementary Table 1.** Information about all MASLD patients used in this study (UCAM and VCU datasets). For each patient we provide the histopathology score (steatosis, ballooning, inflammation, fibrosis, NAS score) and the MASLD stage, as well as additional metadata (age, sex), the Sliding Window (SW) number assigned to the patient, the order in the trajectory (1 to 136), the position according to PC1, the dataset (UCAM or VCU), the batch corrected normalised expression value per gene, and whether or not the gene is present in the gene panels we identify in this study (145 trajectory inference gene set, 57 or 15 biomarker panel).

**Supplementary Table 2.** Patients’ clinical characteristics when divided into CTRL, MASL, MASH F0-1, MASH F2, MASH F3-4. Statistical significance is assessed by ANOVA. The letters next to the values indicate post-hoc analysis significance based on Tukey’s test; “[a]” means reference group; when groups show different letters, they should be considered statistically different at the post-hoc comparison; groups showing the same letter are statistically not-significant at the post-hoc comparison.

**Supplementary Table 3.** Random forest’s accuracy and AUC, tested on UCAM/VCU and GUBRA datasets, using the top 145 genes. The UCAM/VCU dataset has been split into training (80%), and test (20%) sets, and the metrics given in the table are the average over 1000 test-set runs. The gene signature predicts the different MASLD characteristics (Steatosis, Inflammation, Ballooning, Fibrosis) and the MASLD stage. For the GUBRA dataset predictions, the whole UCAM/VCU dataset has been used as a training set. The only available information for this external dataset is the MASLD stage. Finally, for both datasets, a random set of 20, 145, and 5000 genes (averaged across 1000 runs) verifies the robustness of the selected genes.

**Supplementary Table 4.** Pathway enrichment analysis of the 145 genes identified by random forest as key contributors to defining the MASLD disease trajectory.

**Supplementary Table 5.** Patient assignment to each of the 13 Sliding Windows (SWs).

**Supplementary Table 6.** Genes assigned to each module were identified using WGCNA and were associated with the histological phenotypes in our dataset.

**Supplementary Table 7.** Reactome pathway enrichment analysis on the filtered gene sets of each module.

**Supplementary Table 8.** Transcription Factor (TF) enrichment analysis for each significantly-identified gene module.

**Supplementary Table 9.** Reactome pathways enrichment analysis for each MASLD variable-based set of TFs.

**Supplementary Table 10.** Unified inferred global MASLD network according to our analysis. The file includes both the UCAM/Sanyal and EPoS networks and columns E and F indicate whether the edge was present in each of these networks.

**Supplementary Table 11.** Literature-curated genes previously associated with MASLD.

**Supplementary Table 12.** VIPER analysis results comparing consecutive Sliding Windows (SWs). The results are derived from the differentially expressed genes of the previous steps (Log2FoldChange and p-values). The table is organised into 2-plets showing for each comparison the FDR and the Normalised Enrichment Score (NES). NES determines whether a TF has significantly more “activated” than “inhibited” predictions (NES > 0) or vice versa (NES < 0).

**Supplementary Table 13.** Activity scores (cumulative values) of Reactome pathways along the MASLD trajectory for UCAM/VCU and EPoS datasets.

**Supplementary Table 14.** Consensus estimated cell type proportions per sample in the bulk RNA-seq dataset.

**Supplementary Table 15.** FDR values from the enrichment analysis between liver cell type markers and Reactome pathway gene sets, whose differentiation cannot be explained by changes in cell type composition.

**Supplementary Table 16.** GWAS summary results of gene associations to complex traits

**Supplementary Table 17.** Small molecules extracted from the ChEMBL database as targeting one of the proteins represented in our 57-protein biomarker panel.

**Supplementary Table 18.** Fujiwara metadata for all patients with paired biopsies. The table includes Fibrosis scores and NITs (FIB-4, APRI, NFS) scores calculated from the available metadata.

**Supplementary Table 19.** Reference PPI network derived from the integration of multiple interaction databases: KEGG, ReconX, HumanCyc, Reactome, OmniPath. This network was used to generate the global MASLD network.

**Supplementary Table 20.** Slim version of the Reactome database. The hierarchical graph has been pruned using the Information Content (IC) and Semantic Value (SV) properties of terms, retaining the most generic ones. Two columns indicate if a term has been included in the MASLD-related subgraphs of the UCAM-VCU and EpoS datasets

## Notes

### Competing Interest Statement

The authors have declared no competing interest.

### Summary of Updates

The manuscript is heavily revised with additional analyses to confirm and enhance robustness and demonstrate reproducibility of findings, include a set of longitudinal data and improve the biomarker analysis, adding a comparison to state-of-the-art metrics.

## References

1. Chan, W. K. et al. Metabolic Dysfunction-Associated Steatotic Liver Disease (MASLD): A State-of-the-Art Review. Journal of obesity & metabolic syndrome 32, (2023).

2. Loomba, R. & Sanyal, A. J. The global NAFLD epidemic. Nat. Rev. Gastroenterol. Hepatol. 10, (2013).

3. Estes, C. et al. Modeling NAFLD disease burden in China, France, Germany, Italy, Japan, Spain, United Kingdom, and United States for the period 2016-2030. J. Hepatol. 69, (2018).

4. Angulo, P. et al. The NAFLD fibrosis score: a noninvasive system that identifies liver fibrosis in patients with NAFLD. *Hepatology (Baltimore*, Md.) 45, (2007).

5. Sterling, R. K. et al. Development of a simple noninvasive index to predict significant fibrosis in patients with HIV/HCV coinfection. Hepatology 43, 1317–1325 (2006).

6. Kleiner, D. E. et al. Design and validation of a histological scoring system for nonalcoholic fatty liver disease. Hepatology 41, 1313–1321 (2005).

7. McGill, D. B., Rakela, J., Zinsmeister, A. R. & Ott, B. J. A 21-year experience with major hemorrhage after percutaneous liver biopsy. Gastroenterology 99, (1990).

8. Ratziu, V. et al. Sampling variability of liver biopsy in nonalcoholic fatty liver disease. Gastroenterology 128, (2005).

9. Chalasani, N. et al. The diagnosis and management of nonalcoholic fatty liver disease: Practice guidance from the American Association for the Study of Liver Diseases. *Hepatology (Baltimore*, Md.) 67, (2018).

10. EASL-EASD-EASO Clinical Practice Guidelines on the management of metabolic dysfunction-associated steatotic liver disease (MASLD). Journal of hepatology 81, (2024).

11. Castera, L., Forns, X. & Alberti, A. Non-invasive evaluation of liver fibrosis using transient elastography. Journal of hepatology 48, (2008).

12. Bedossa, P. Utility and appropriateness of the fatty liver inhibition of progression (FLIP) algorithm and steatosis, activity, and fibrosis (SAF) score in the evaluation of biopsies of nonalcoholic fatty liver disease. *Hepatology (Baltimore*, Md.) 60, (2014).

13. Govaere, O. et al. Transcriptomic profiling across the nonalcoholic fatty liver disease spectrum reveals gene signatures for steatohepatitis and fibrosis. Sci. Transl. Med. 12, (2020).

14. Hoang, S. A. et al. Gene Expression Predicts Histological Severity and Reveals Distinct Molecular Profiles of Nonalcoholic Fatty Liver Disease. Sci. Rep. 9, 12541 (2019).

15. Atabaki-Pasdar, N. et al. Predicting and elucidating the etiology of fatty liver disease: A machine learning modeling and validation study in the IMI DIRECT cohorts. PLoS Med. 17, e1003149 (2020).

16. Anstee, Q. M., Seth, D. & Day, C. P. Genetic Factors That Affect Risk of Alcoholic and Nonalcoholic Fatty Liver Disease. Gastroenterology 150, (2016).

17. Wong, V. W.-S., Adams, L. A., de Lédinghen, V., Wong, G. L.-H. & Sookoian, S. Noninvasive biomarkers in NAFLD and NASH - current progress and future promise. Nat. Rev. Gastroenterol. Hepatol. 15, 461–478 (2018).

18. Eslam, M. & George, J. Genetic contributions to NAFLD: leveraging shared genetics to uncover systems biology. Nat. Rev. Gastroenterol. Hepatol. 17, 40–52 (2019).

19. Chen, Y. et al. Genome-wide association meta-analysis identifies 17 loci associated with nonalcoholic fatty liver disease. Nat. Genet. 55, 1640–1650 (2023).

20. Fairfield, C. J. et al. Genome-Wide Association Study of NAFLD Using Electronic Health Records. Hepatology Communications 6, 297 (2022).

21. Genome-wide association study of non-alcoholic fatty liver and steatohepatitis in a histologically characterised cohort⋆. J. Hepatol. 73, 505–515 (2020).

22. Romeo, S. et al. Genetic variation in PNPLA3 confers susceptibility to nonalcoholic fatty liver disease. Nature Genetics 40, 1461–1465 (2008).

23. Rotman, Y., Koh, C., Zmuda, J. M., Kleiner, D. E. & Liang, T. J. The association of genetic variability in patatin-like phospholipase domain-containing protein 3 (PNPLA3) with histological severity of nonalcoholic fatty liver disease. *Hepatology (Baltimore*, Md.) 52, (2010).

24. Kozlitina, J. et al. Exome-wide association study identifies a TM6SF2 variant that confers susceptibility to nonalcoholic fatty liver disease. Nature genetics 46, (2014).

25. Mancina, R. M. et al. The MBOAT7-TMC4 Variant rs641738 Increases Risk of Nonalcoholic Fatty Liver Disease in Individuals of European Descent. Gastroenterology 150, (2016).

26. Abul-Husn, N. S. et al. A Protein-Truncating HSD17B13 Variant and Protection from Chronic Liver Disease. New England Journal of Medicine (2018) doi:10.1056/NEJMoa1712191.

27. Magwene, P. M., Lizardi, P. & Kim, J. Reconstructing the temporal ordering of biological samples using microarray data. Bioinformatics 19, 842–850 (2003).

28. Mukherjee, S. et al. Molecular estimation of neurodegeneration pseudotime in older brains. Nat. Commun. 11, 5781 (2020).

29. Young, A. L. et al. Data-driven modelling of neurodegenerative disease progression: thinking outside the black box. Nat. Rev. Neurosci. 25, 111–130 (2024).

30. Cherubini, A., Della Torre, S., Pelusi, S. & Valenti, L. Sexual dimorphism of metabolic dysfunction-associated steatotic liver disease. Trends in molecular medicine 30, (2024).

31. Glass, L. M., Hunt, C. M., Fuchs, M. & Su, G. L. Comorbidities and Nonalcoholic Fatty Liver Disease: The Chicken, the Egg, or Both? Federal Practitioner 36, 64 (2019).

32. Rosato, V. et al. NAFLD and Extra-Hepatic Comorbidities: Current Evidence on a Multi-Organ Metabolic Syndrome. International Journal of Environmental Research and Public Health 16, 3415 (2019).

33. Azzu, V. et al. Suppression of insulin-induced gene 1 (INSIG1) function promotes hepatic lipid remodelling and restrains NASH progression. Molecular metabolism 48, (2021).

34. Suppli, M. P. et al. Hepatic transcriptome signatures in patients with varying degrees of nonalcoholic fatty liver disease compared with healthy normal-weight individuals. Am. J. Physiol. Gastrointest. Liver Physiol. 316, (2019).

35. Fujiwara, N. et al. Molecular signatures of long-term hepatocellular carcinoma risk in nonalcoholic fatty liver disease. Science translational medicine 14, (2022).

36. Barker, C. G. et al. Identification of phenotype-specific networks from paired gene expression–cell shape imaging data. Genome Res. 32, 750 (2022).

37. Langfelder, P. & Horvath, S. WGCNA: an R package for weighted correlation network analysis. BMC Bioinformatics 9, 1–13 (2008).

38. Milacic, M. et al. The Reactome Pathway Knowledgebase 2024. Nucleic Acids Res. 52, D672–D678 (2023).

39. Maiers, J. L. & Malhi, H. Endoplasmic Reticulum Stress in Metabolic Liver Diseases and Hepatic Fibrosis. Seminars in liver disease 39, (2019).

40. Ortiz, C. et al. Extracellular Matrix Remodeling in Chronic Liver Disease. Current tissue microenvironment reports 2, (2021).

41. Nassir, F. NAFLD: Mechanisms, Treatments, and Biomarkers. Biomolecules 12, (2022).

42. Córdova, G. et al. SMAD3 and SP1/SP3 Transcription Factors Collaborate to Regulate Connective Tissue Growth Factor Gene Expression in Myoblasts in Response to Transforming Growth Factor β. J. Cell. Biochem. 116, 1880–1887 (2015).

43. Younesi, F. S., Miller, A. E., Barker, T. H., Rossi, F. M. V. & Hinz, B. Fibroblast and myofibroblast activation in normal tissue repair and fibrosis. Nat. Rev. Mol. Cell Biol. 25, 617–638 (2024).

44. Liu, D. et al. Ets-1 deficiency alleviates nonalcoholic steatohepatitis via weakening TGF-β1 signaling-mediated hepatocyte apoptosis. Cell Death Dis. 10, 1–14 (2019).

45. Varga, T., Czimmerer, Z. & Nagy, L. PPARs are a unique set of fatty acid regulated transcription factors controlling both lipid metabolism and inflammation. Biochim. Biophys. Acta 1812, 1007 (2011).

46. Vachliotis, I. D. & Polyzos, S. A. The Role of Tumor Necrosis Factor-Alpha in the Pathogenesis and Treatment of Nonalcoholic Fatty Liver Disease. Curr. Obes. Rep. 12, 191–206 (2023).

47. Tumor necrosis factor-α signaling in nonalcoholic steatohepatitis and targeted therapies. J. Genet. Genomics 49, 269–278 (2022).

48. Parviz, F. et al. Hepatocyte nuclear factor 4α controls the development of a hepatic epithelium and liver morphogenesis. Nature Genetics 34, 292–296 (2003).

49. Tontonoz, P. & Spiegelman, B. M. Fat and beyond: the diverse biology of PPARgamma. Annual review of biochemistry 77, (2008).

50. Bitter, A. et al. Human sterol regulatory element-binding protein 1a contributes significantly to hepatic lipogenic gene expression. Cellular physiology and biochemistry: international journal of experimental cellular physiology, biochemistry, and pharmacology 35, (2015).

51. Luedde, T. & Schwabe, R. F. NF-κB in the liver—linking injury, fibrosis and hepatocellular carcinoma. Nat. Rev. Gastroenterol. Hepatol. 8, 108 (2011).

52. Luo, M., Li, T. & Sang, H. The role of hypoxia-inducible factor 1α in hepatic lipid metabolism. J. Mol. Med. 101, 487–500 (2023).

53. Mesarwi, O. A. et al. Hepatocyte Hypoxia Inducible Factor-1 Mediates the Development of Liver Fibrosis in a Mouse Model of Nonalcoholic Fatty Liver Disease. PLoS One 11, e0168572 (2016).

54. CREB family: A significant role in liver fibrosis. Biochimie 163, 94–100 (2019).

55. The Molecular Signatures Database Hallmark Gene Set Collection. Cell Systems 1, 417–425 (2015).

56. Browning, J. D. & Horton, J. D. Molecular mediators of hepatic steatosis and liver injury. J Clin Invest 114, 147–152 (2004).

57. Nagaya, T. et al. Down-regulation of SREBP-1c is associated with the development of burned-out NASH. Journal of hepatology 53, (2010).

58. Xu, Y. et al. Hepatocyte Nuclear Factor 4α Prevents the Steatosis-to-NASH Progression by Regulating p53 and Bile Acid Signaling (in mice). Hepatology 73, 2251–2265 (2021).

59. Maglich, J. M., Lobe, D. C. & Moore, J. T. The nuclear receptor CAR (NR1I3) regulates serum triglyceride levels under conditions of metabolic stress. Journal of lipid research 50, (2009).

60. Tan, J. et al. HNF1α Controls Liver Lipid Metabolism and Insulin Resistance via Negatively Regulating the SOCS-3-STAT3 Signaling Pathway. Journal of Diabetes Research 2019, 5483946 (2019).

61. Zhang, S. et al. Smad7 Antagonizes Transforming Growth Factor β Signaling in the Nucleus by Interfering with Functional Smad-DNA Complex Formation. Molecular and Cellular Biology 27, 4488 (2007).

62. Lu, J., McKinsey, T. A., Zhang, C. L. & Olson, E. N. Regulation of skeletal myogenesis by association of the MEF2 transcription factor with class II histone deacetylases. Mol. Cell 6, (2000).

63. Mann, D. A. & Smart, D. E. Transcriptional regulation of hepatic stellate cell activation. Gut 50, 891 (2002).

64. Schüler, A. et al. The MADS transcription factor Mef2c is a pivotal modulator of myeloid cell fate. Blood 111, 4532–4541 (2008).

65. Cilenti, F. et al. A PGE2-MEF2A axis enables context-dependent control of inflammatory gene expression. Immunity 54, (2021).

66. Liu, N. et al. Notch and retinoic acid signals regulate macrophage formation from endocardium downstream of Nkx2-5. Nature Communications 14, 5398 (2023).

67. Mu, W. et al. Potential Nexus of Non-alcoholic Fatty Liver Disease and Type 2 Diabetes Mellitus: Insulin Resistance Between Hepatic and Peripheral Tissues. Front. Pharmacol. 9, 430129 (2019).

68. Guilliams, M. et al. Spatial proteogenomics reveals distinct and evolutionarily conserved hepatic macrophage niches. Cell 185, (2022).

69. Govaere, O. et al. A proteo-transcriptomic map of non-alcoholic fatty liver disease signatures. Nature Metabolism 5, 572 (2023).

70. Buniello, A. et al. The NHGRI-EBI GWAS Catalog of published genome-wide association studies, targeted arrays and summary statistics 2019. Nucleic acids research 47, (2019).

71. Chitturi, S. et al. The Asia-Pacific Working Party on Non-alcoholic Fatty Liver Disease guidelines 2017-Part 2: Management and special groups. J. Gastroenterol. Hepatol. 33, (2018).

72. Wnt/beta-catenin signaling and its modulators in nonalcoholic fatty liver diseases. Hepatobiliary Pancreat. Dis. Int 22, 333–345 (2023).

73. Ying, H.-Z. et al. PDGF signaling pathway in hepatic fibrosis pathogenesis and therapeutics. Mol. Med. Rep. 16, 7879 (2017).

74. TGF-β1 signaling can worsen NAFLD with liver fibrosis backdrop. Exp. Mol. Pathol. 124, 104733 (2022).

75. Gribben, C. et al. Acquisition of epithelial plasticity in human chronic liver disease. Nature 630, 166–173 (2024).

76. Vonderlin, J., Chavakis, T., Sieweke, M. & Tacke, F. The Multifaceted Roles of Macrophages in NAFLD Pathogenesis. Cell Mol Gastroenterol Hepatol 15, 1311–1324 (2023).

77. Matchett, K. P., Paris, J., Teichmann, S. A. & Henderson, N. C. Spatial genomics: mapping human steatotic liver disease. Nat. Rev. Gastroenterol. Hepatol. 1–15 (2024).

78. Harrison, S. A. et al. A blood-based biomarker panel (NIS4) for non-invasive diagnosis of non-alcoholic steatohepatitis and liver fibrosis: a prospective derivation and global validation study. *The lancet*. Gastroenterology & hepatology 5, (2020).

79. Levy, M. T., McCaughan, G. W., Marinos, G. & Gorrell. Intrahepatic expression of the hepatic stellate cell marker fibroblast activation protein correlates with the degree of fibrosis in hepatitis C virus infection. Liver 22, (2002).

80. Yang, A. T. et al. Fibroblast Activation Protein Activates Macrophages and Promotes Parenchymal Liver Inflammation and Fibrosis. Cellular and molecular gastroenterology and hepatology 15, (2023).

81. Kozumi, K. et al. Transcriptomics Identify Thrombospondin-2 as a Biomarker for NASH and Advanced Liver Fibrosis. *Hepatology (Baltimore*, Md.) 74, (2021).

82. Lee, C. H. et al. Circulating Thrombospondin-2 as a Novel Fibrosis Biomarker of Nonalcoholic Fatty Liver Disease in Type 2 Diabetes. Diabetes care 44, (2021).

83. Friedman, S. L., Neuschwander-Tetri, B. A., Rinella, M. & Sanyal, A. J. Mechanisms of NAFLD development and therapeutic strategies. Nat. Med. 24, 908–922 (2018).

84. Heebøll, S. et al. Placebo-controlled, randomised clinical trial: high-dose resveratrol treatment for non-alcoholic fatty liver disease. Scand. J. Gastroenterol. (2016) doi:10.3109/00365521.2015.1107620.

85. Vacca, M. et al. An unbiased ranking of murine dietary models based on their proximity to human metabolic dysfunction-associated steatotic liver disease (MASLD). Nature Metabolism 6, 1178–1196 (2024).

86. Wai, C. T. et al. A simple noninvasive index can predict both significant fibrosis and cirrhosis in patients with chronic hepatitis C. *Hepatology (Baltimore*, Md.) 38, (2003).

87. Kim, D., Langmead, B. & Salzberg, S. L. HISAT: a fast spliced aligner with low memory requirements. Nat. Methods 12, 357–360 (2015).

88. Anders, S., Pyl, P. T. & Huber, W. HTSeq—a Python framework to work with high-throughput sequencing data. Bioinformatics 31, 166–169 (2014).

89. Durinck, S., Spellman, P. T., Birney, E. & Huber, W. Mapping identifiers for the integration of genomic datasets with the R/Bioconductor package biomaRt. Nat. Protoc. 4, (2009).

90. Zhang, Y., Jenkins, D. F., Manimaran, S. & Johnson, W. E. Alternative empirical Bayes models for adjusting for batch effects in genomic studies. BMC Bioinformatics 19, 1–15 (2018).

91. Saelens, W., Cannoodt, R., Todorov, H. & Saeys, Y. A comparison of single-cell trajectory inference methods. Nat. Biotechnol. 37, 547–554 (2019).

92. Street, K. et al. Slingshot: cell lineage and pseudotime inference for single-cell transcriptomics. BMC Genomics 19, 1–16 (2018).

93. Fisher, R. A. Statistical Methods for Research Workers. Breakthroughs in Statistics 66–70 (1992).

94. Benjamini, Y. & Hochberg, Y. Controlling the False Discovery Rate: A Practical and Powerful Approach to Multiple Testing. J. R. Stat. Soc. Series B Stat. Methodol. 57, 289–300 (1995).

95. Zhang, B. & Horvath, S. A General Framework for Weighted Gene Co-Expression Network Analysis. Stat. Appl. Genet. Mol. Biol. 4, (2005).

96. Stoica, P. & Selen, Y. Model-order selection: a review of information criterion rules. https://ieeexplore.ieee.org/document/1311138.

97. Alvarez, M. J. et al. Network-based inference of protein activity helps functionalize the genetic landscape of cancer. Nat. Genet. 48, 838 (2016).

98. Müller-Dott, S. et al. Expanding the coverage of regulons from high-confidence prior knowledge for accurate estimation of transcription factor activities. Nucleic Acids Res 51, 10934–10949 (2023).

99. Akhmedov, M. et al. PCSF: An R-package for network-based interpretation of high-throughput data. PLoS Comput. Biol. 13, e1005694 (2017).

100. Kuleshov, M. V. et al. Enrichr: a comprehensive gene set enrichment analysis web server 2016 update. Nucleic Acids Res. 44, W90–W97 (2016).

101. Lachmann, A. et al. ChEA: transcription factor regulation inferred from integrating genome-wide ChIP-X experiments. Bioinformatics 26, 2438–2444 (2010).

102. Stormo, G. D. Modeling the specificity of protein-DNA interactions. Quantitative Biology 1, 115–130 (2013).

103. Wasserman, W. W. & Sandelin, A. Applied bioinformatics for the identification of regulatory elements. Nat. Rev. Genet. 5, 276–287 (2004).

104. Gillespie, M. et al. The reactome pathway knowledgebase 2022. Nucleic Acids Res. 50, D687–D692 (2021).

105. Kanehisa, M. & Goto, S. KEGG: kyoto encyclopedia of genes and genomes. Nucleic Acids Res. 28, (2000).

106. Wrzodek, C., Büchel, F., Ruff, M., Dräger, A. & Zell, A. Precise generation of systems biology models from KEGG pathways. BMC Syst. Biol. 7, (2013).

107. Romero, P. et al. Computational prediction of human metabolic pathways from the complete human genome. Genome Biol. 6, (2005).

108. Thiele, I. et al. A community-driven global reconstruction of human metabolism. Nat. Biotechnol. 31, (2013).

109. Rodchenkov, I. et al. Pathway Commons 2019 Update: integration, analysis and exploration of pathway data. Nucleic Acids Res. 48, D489–D497 (2019).

110. Türei, D., Korcsmáros, T. & Saez-Rodriguez, J. OmniPath: guidelines and gateway for literature-curated signaling pathway resources. Nat. Methods 13, 966–967 (2016).

111. Türei, D. et al. Integrated intra-and intercellular signaling knowledge for multicellular omics analysis. Mol. Syst. Biol. 17, e9923 (2021).

112. OmnipathR. Bioconductor http://bioconductor.org/packages/OmnipathR/.

113. Csárdi, G., et al. Igraph: Network analysis and visualization. CRAN: Contributed Packages The R Foundation 10.32614/cran.package.igraph (2006).

114. Aleksander, S. A. et al. The Gene Ontology knowledgebase in 2023. Genetics 224, (2023).

115. Yu, G. et al. GOSemSim: an R package for measuring semantic similarity among GO terms and gene products. Bioinformatics 26, 976–978 (2010).

116. Resnik, P. Semantic Similarity in a Taxonomy: An Information-Based Measure and its Application to Problems of Ambiguity in Natural Language. jair 11, 95–130 (1999).

117. An Information-Theoretic Definition of Similarity. doi:10.5555/645527.657297.

118. Wang, J. Z., Du, Z., Payattakool, R., Yu, P. S. & Chen, C.-F. A new method to measure the semantic similarity of GO terms. Bioinformatics 23, 1274–1281 (2007).

119. Charmpi, K., Chokkalingam, M., Johnen, R. & Beyer, A. Optimizing network propagation for multi-omics data integration. PLoS Comput. Biol. 17, e1009161 (2021).

120. En masse organoid phenotyping informs metabolic-associated genetic susceptibility to NASH. Cell 185, 4216–4232.e16 (2022).

121. Graupera, I. et al. LiverScreen project: study protocol for screening for liver fibrosis in the general population in European countries. BMC Public Health 22, 1–10 (2022).

122. Hendriks, D. et al. Engineered human hepatocyte organoids enable CRISPR-based target discovery and drug screening for steatosis. Nat. Biotechnol. 41, 1567–1581 (2023).

123. Lee, E., Korf, H. & Vidal-Puig, A. An adipocentric perspective on the development and progression of non-alcoholic fatty liver disease. Journal of hepatology 78, (2023).

124. A network-based computational and experimental framework for repurposing compounds toward the treatment of non-alcoholic fatty liver disease. iScience 25, 103890 (2022).

125. Love, M. I., Huber, W. & Anders, S. Moderated estimation of fold change and dispersion for RNA-seq data with DESeq2. Genome Biol. 15, 1–21 (2014).

126. The anatomy of a large-scale hypertextual Web search engine. Computer Networks and ISDN Systems 30, 107–117 (1998).

127. Measure the semantic similarity of GO terms using aggregate information content. IEEE/ACM Transactions on Computational Biology and Bioinformatics (TCBB) (2014) doi:10.1109/TCBB.2013.176.

128. Hao, Y. et al. Dictionary learning for integrative, multimodal and scalable single-cell analysis. Nature Biotechnology 42, 293–304 (2023).

129. Vathrakokoili, P. A. et al. CATD: a reproducible pipeline for selecting cell-type deconvolution methods across tissues. Bioinformatics advances 4, (2024).

130. Zheng, S. C. et al. EpiDISH web server: Epigenetic Dissection of Intra-Sample-Heterogeneity with online GUI. *Bioinformatics (Oxford*, England) 36, (2019).

131. Baron, M. et al. A Single-Cell Transcriptomic Map of the Human and Mouse Pancreas Reveals Inter-and Intra-cell Population Structure. Cell systems 3, (2016).

132. Tsoucas, D. et al. Accurate estimation of cell-type composition from gene expression data. Nature Communications 10, 1–9 (2019).

133. Newman, A. M. et al. Robust enumeration of cell subsets from tissue expression profiles. Nature Methods 12, 453–457 (2015).

134. Hao, Y., Yan, M., Heath, B. R., Lei, Y. L. & Xie, Y. Fast and robust deconvolution of tumor infiltrating lymphocyte from expression profiles using least trimmed squares. PLoS computational biology 15, (2019).

135. Chu, T., Wang, Z., Pe’er, D. & Danko, C. G. Cell type and gene expression deconvolution with BayesPrism enables Bayesian integrative analysis across bulk and single-cell RNA sequencing in oncology. Nature Cancer 3, 505–517 (2022).

136. Satija, R., Farrell, J. A., Gennert, D., Schier, A. F. & Regev, A. Spatial reconstruction of single-cell gene expression data. Nature Biotechnology 33, 495–502 (2015).

137. Mendez, D. et al. ChEMBL: towards direct deposition of bioassay data. Nucleic acids research 47, (2019).

138. Yilmaz, Y. & Eren, F. Serum biomarkers of fibrosis and extracellular matrix remodeling in patients with nonalcoholic fatty liver disease: association with liver histology. European journal of gastroenterology & hepatology 31, (2019).

139. Coilly, A., Desterke, C., Guettier, C., Samuel, D. & Chiappini, F. FABP4 and MMP9 levels identified as predictive factors for poor prognosis in patients with nonalcoholic fatty liver using data mining approaches and gene expression analysis. Scientific reports 9, (2019).

140. Toyoda, H. et al. Higher hepatic gene expression and serum levels of matrix metalloproteinase-2 are associated with steatohepatitis in non-alcoholic fatty liver diseases. Biomarkers: biochemical indicators of exposure, response, and susceptibility to chemicals 18, (2013).

141. Qi, S., Wang, C., Li, C., Wang, P. & Liu, M. Candidate genes investigation for severe nonalcoholic fatty liver disease based on bioinformatics analysis. Medicine 96, (2017).

