## Supplementary figures and images for "Decoding MASLD Progression: A Molecular Trajectory-Based Framework for Modelling Disease Dynamics"

### Figure S1

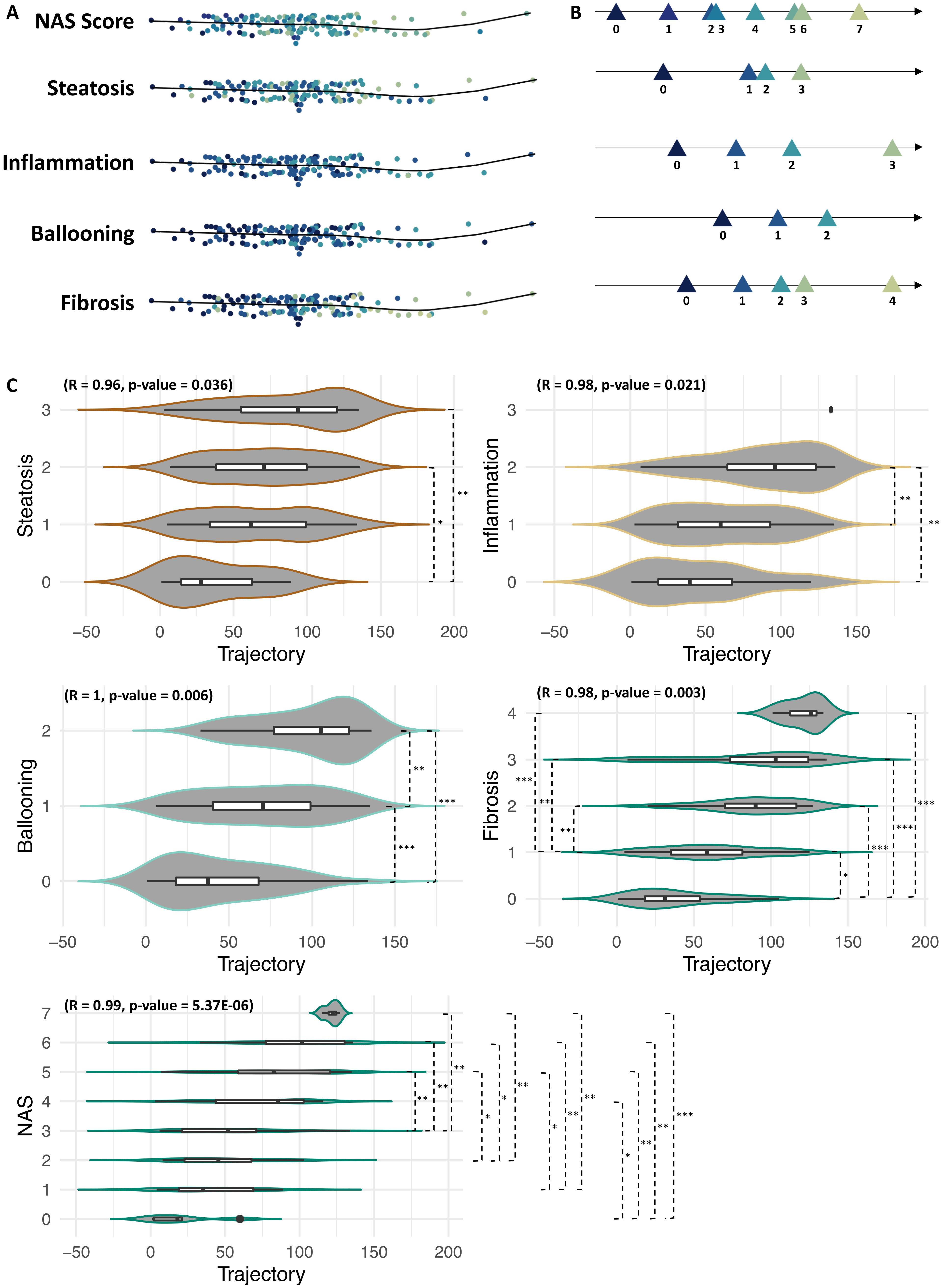

### Figure S2

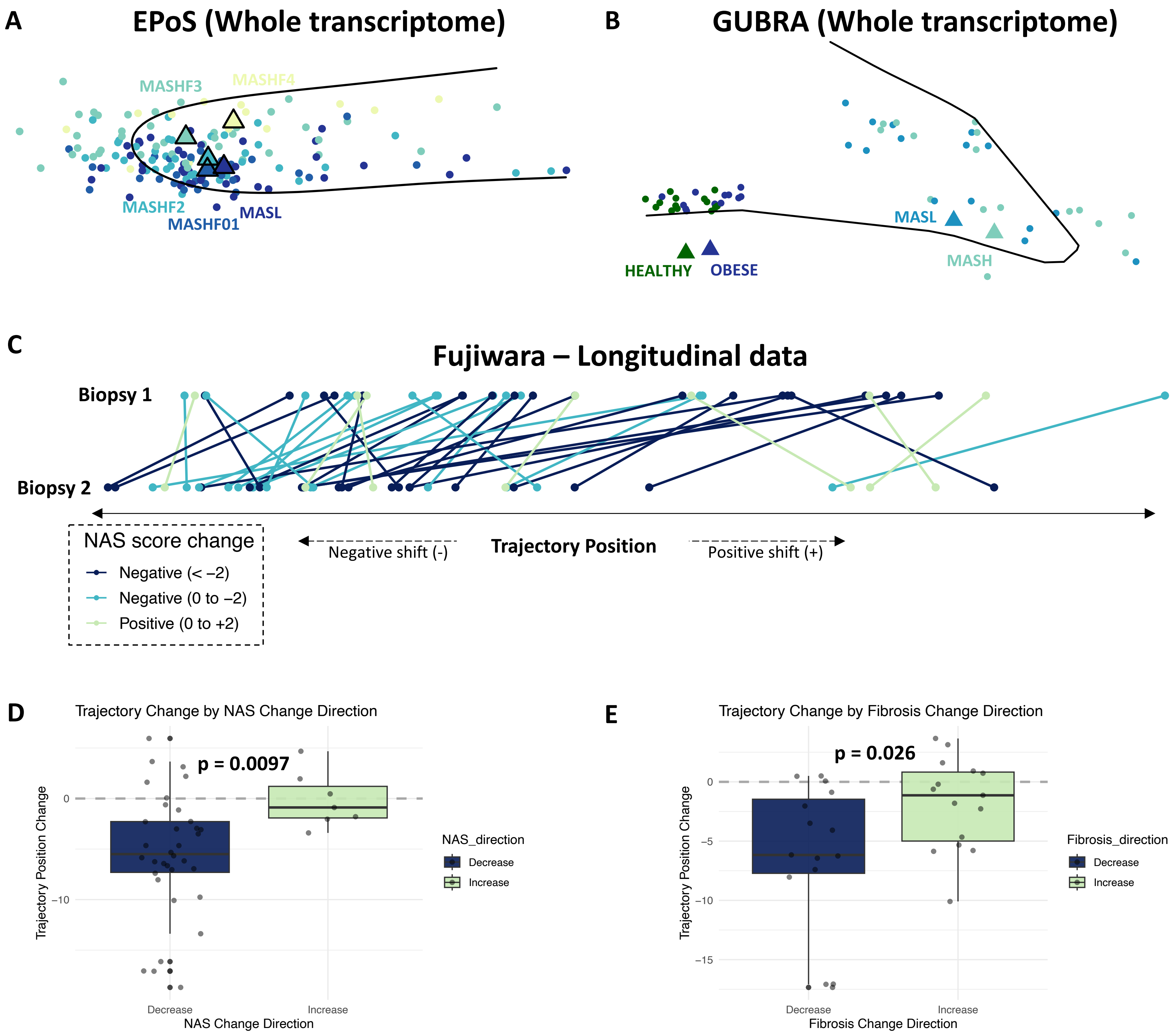

### Figure S3

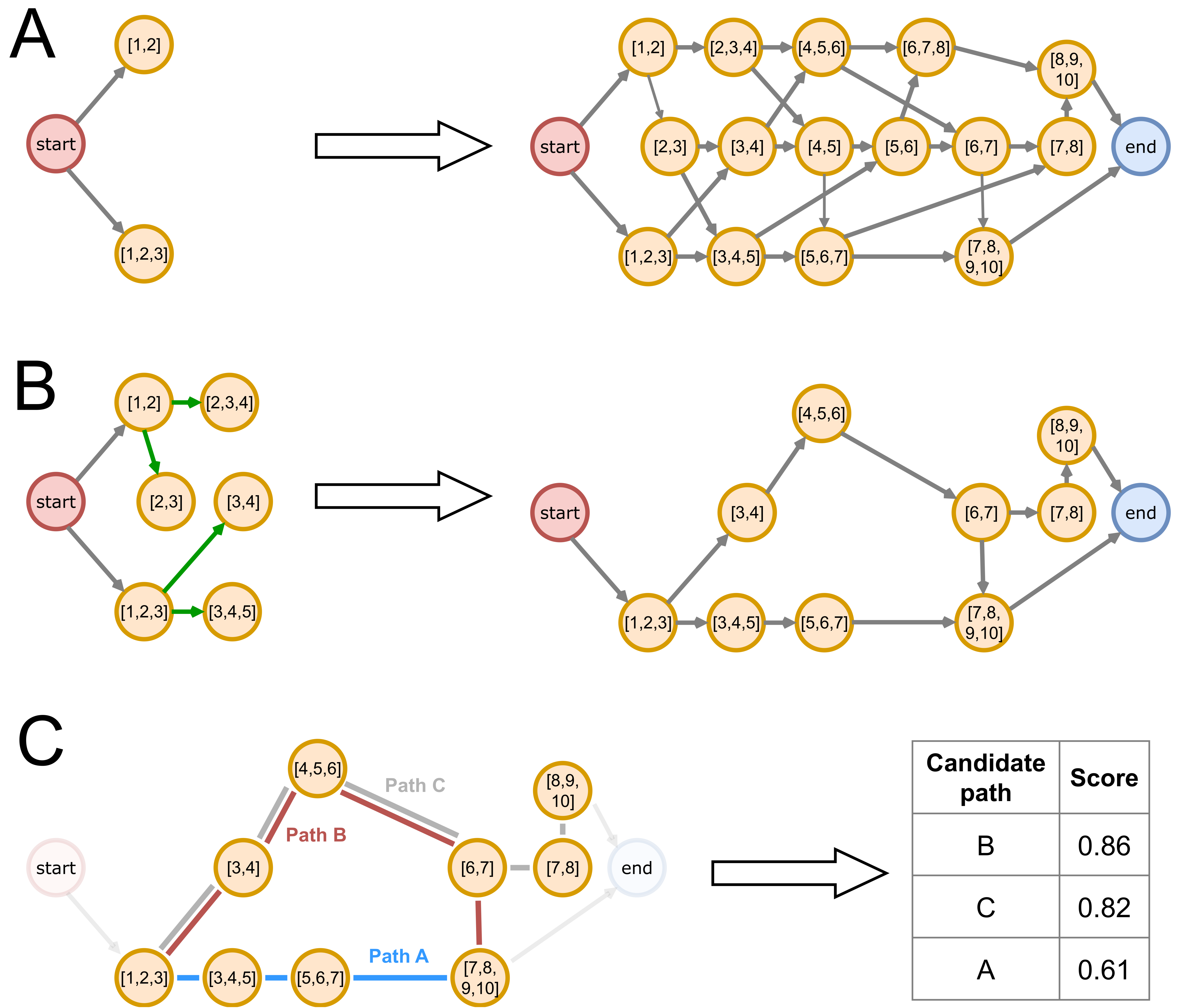


| Candidate path | Score |
|---|---|
| B | 0.86 |
| C | 0.82 |
| A | 0.61 |

### Figure S4

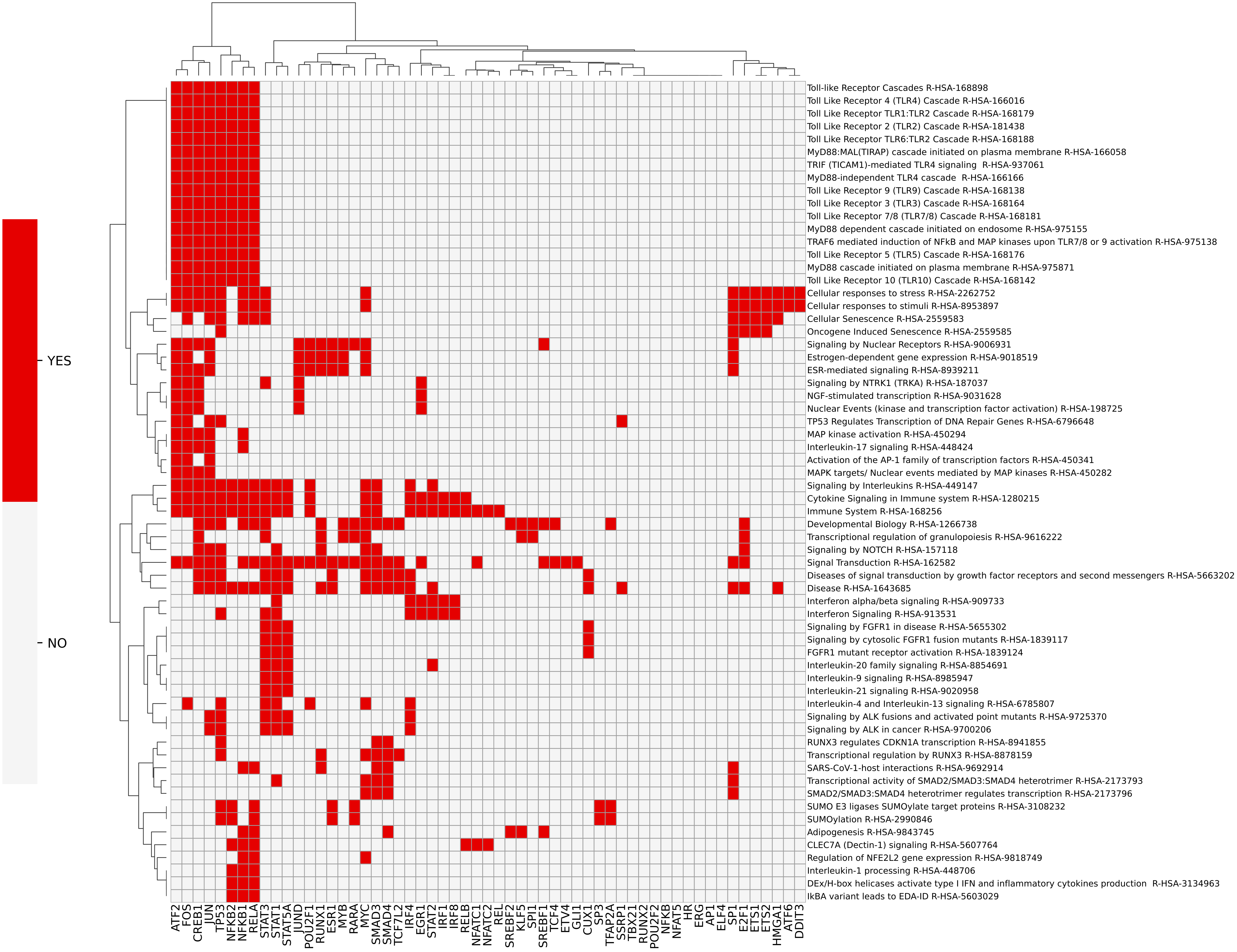

### Figure S5

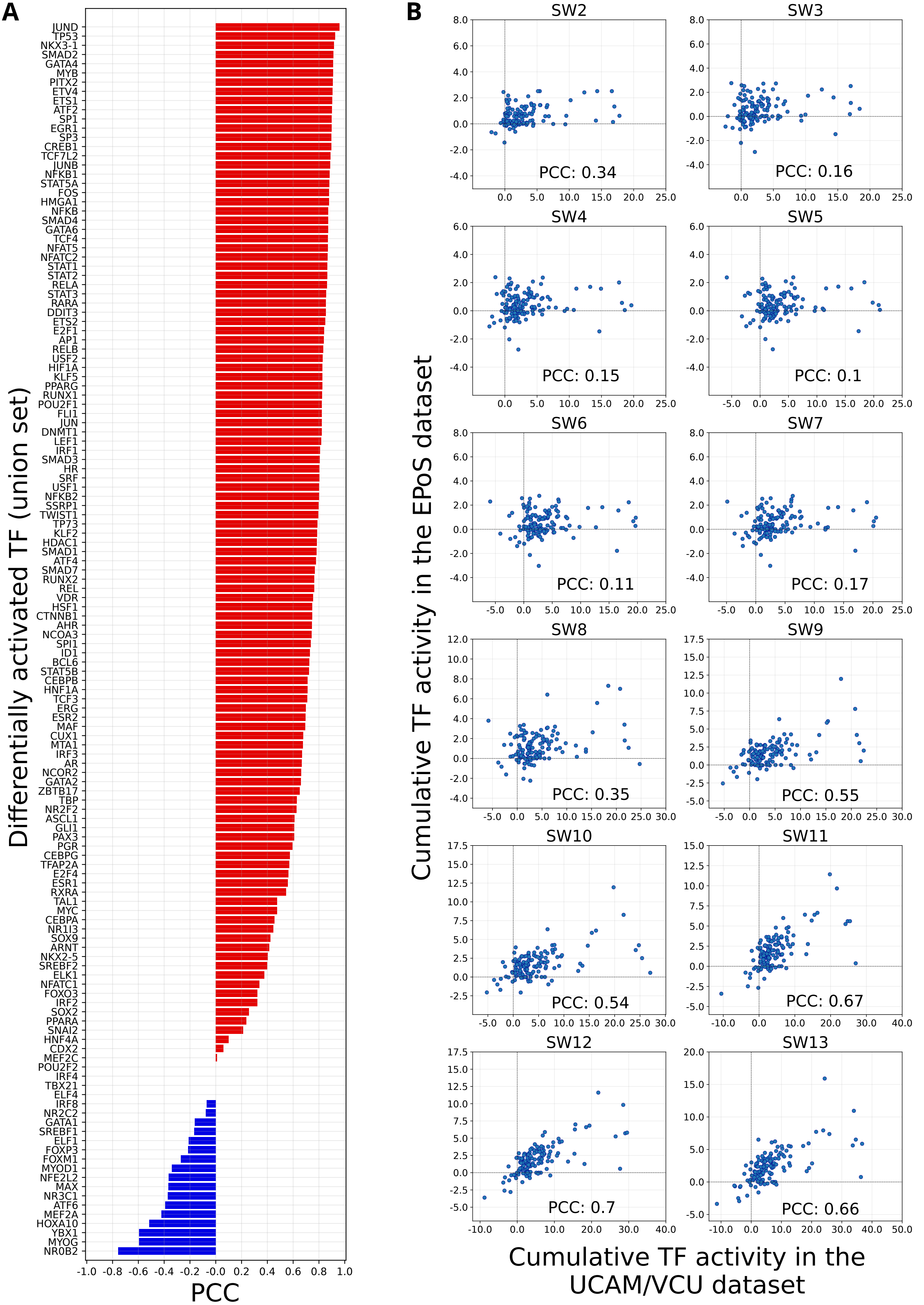

### Figure S6

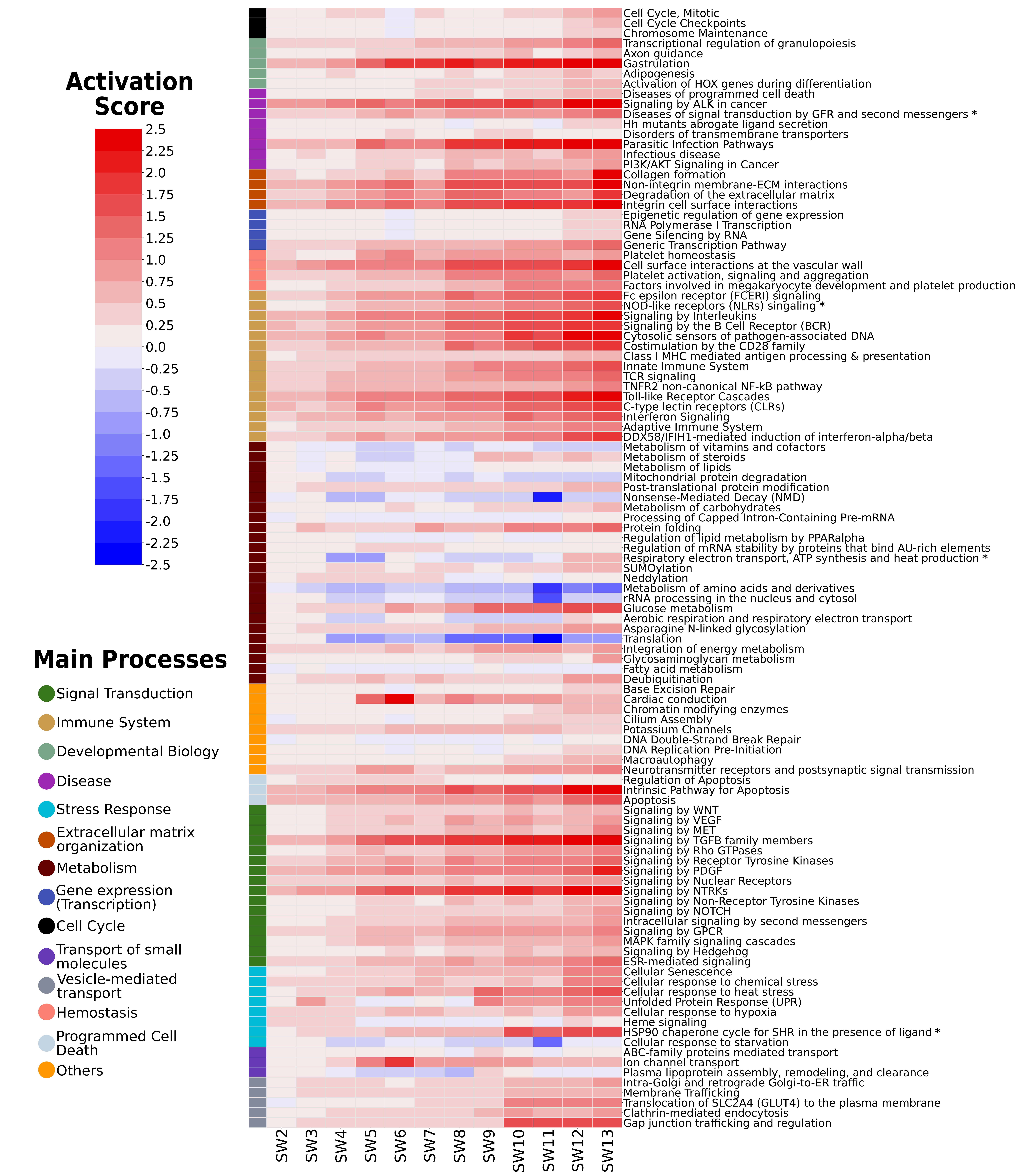

### Figure S7

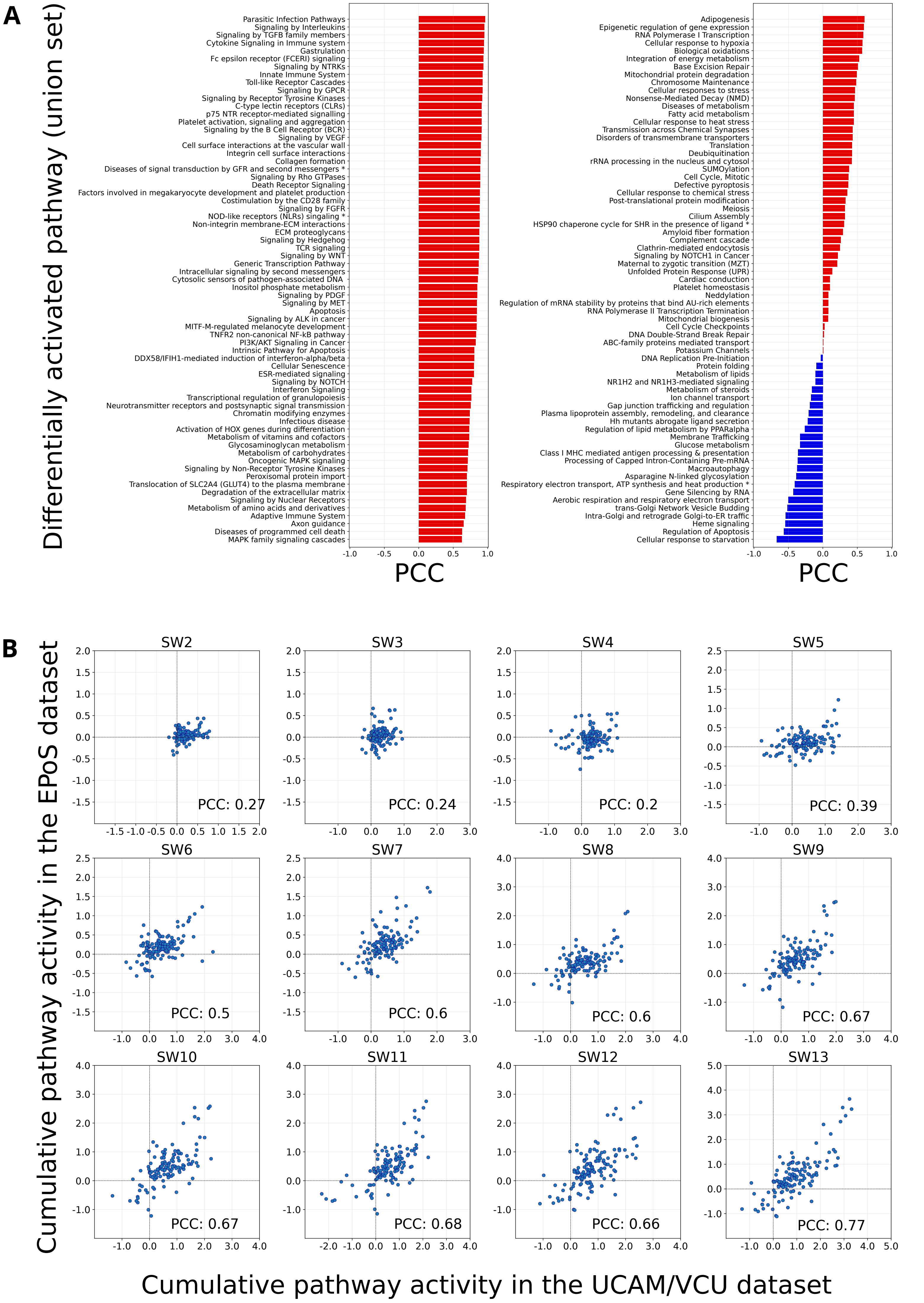

### Figure S8

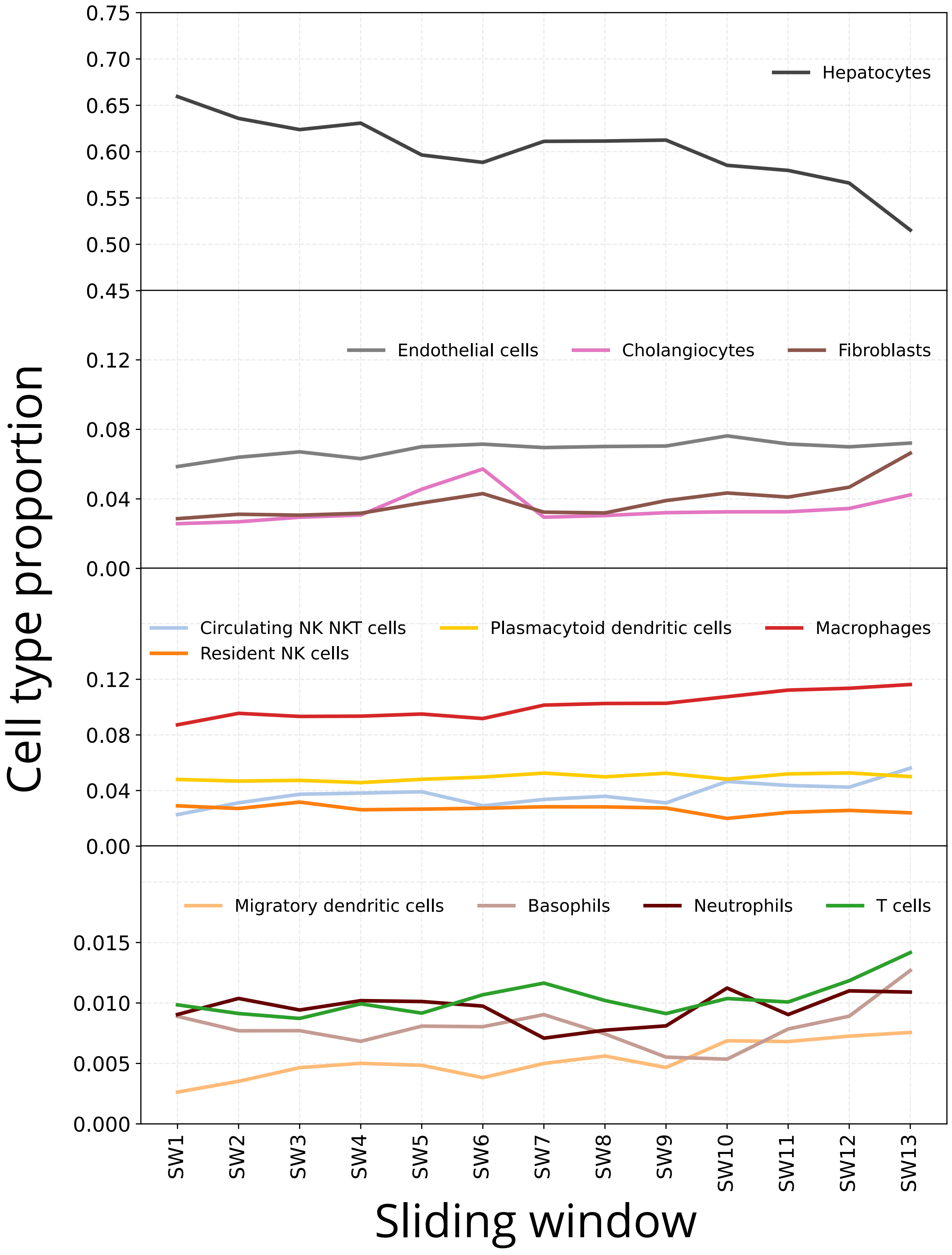

### Figure S9

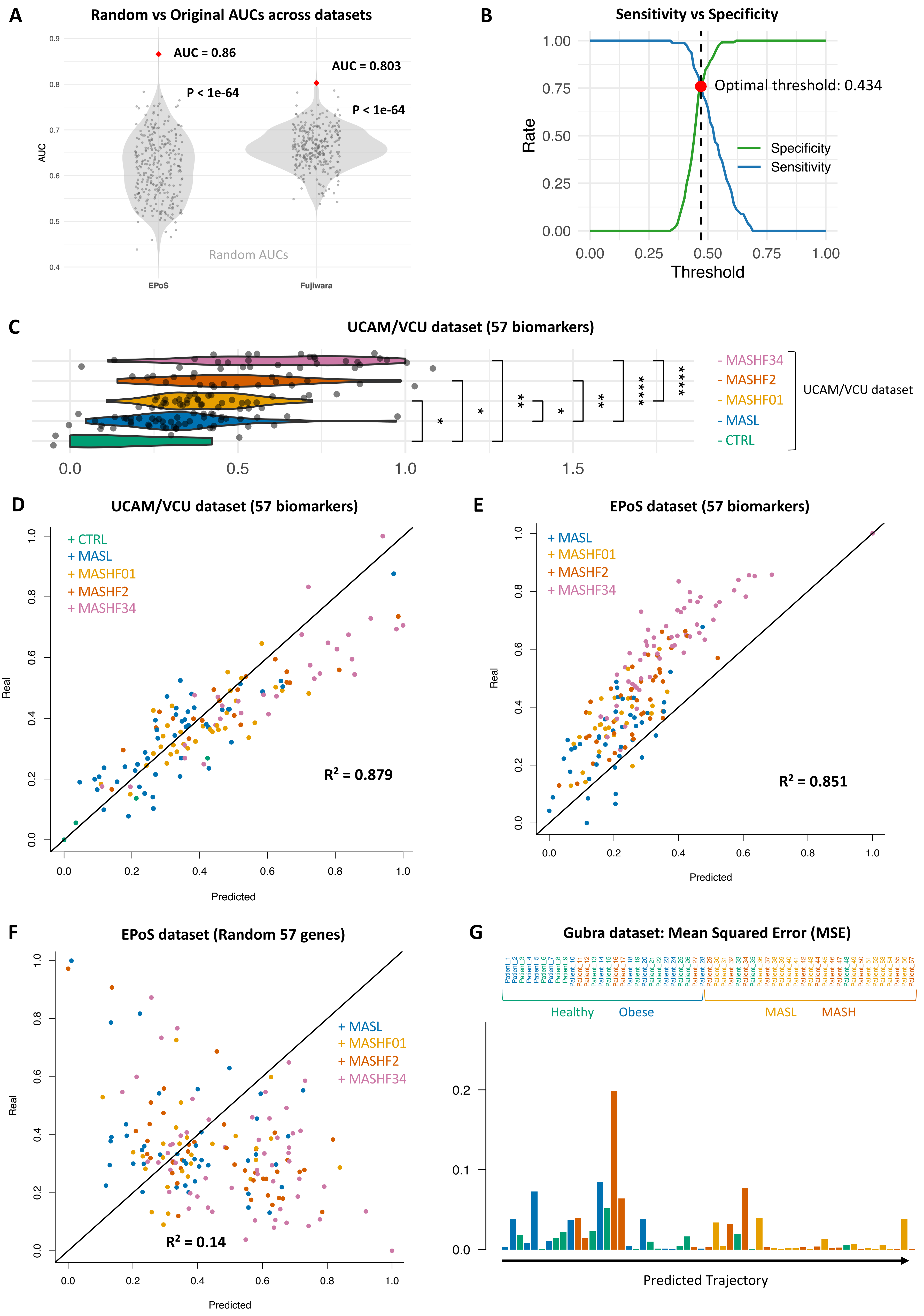

### Figure S10

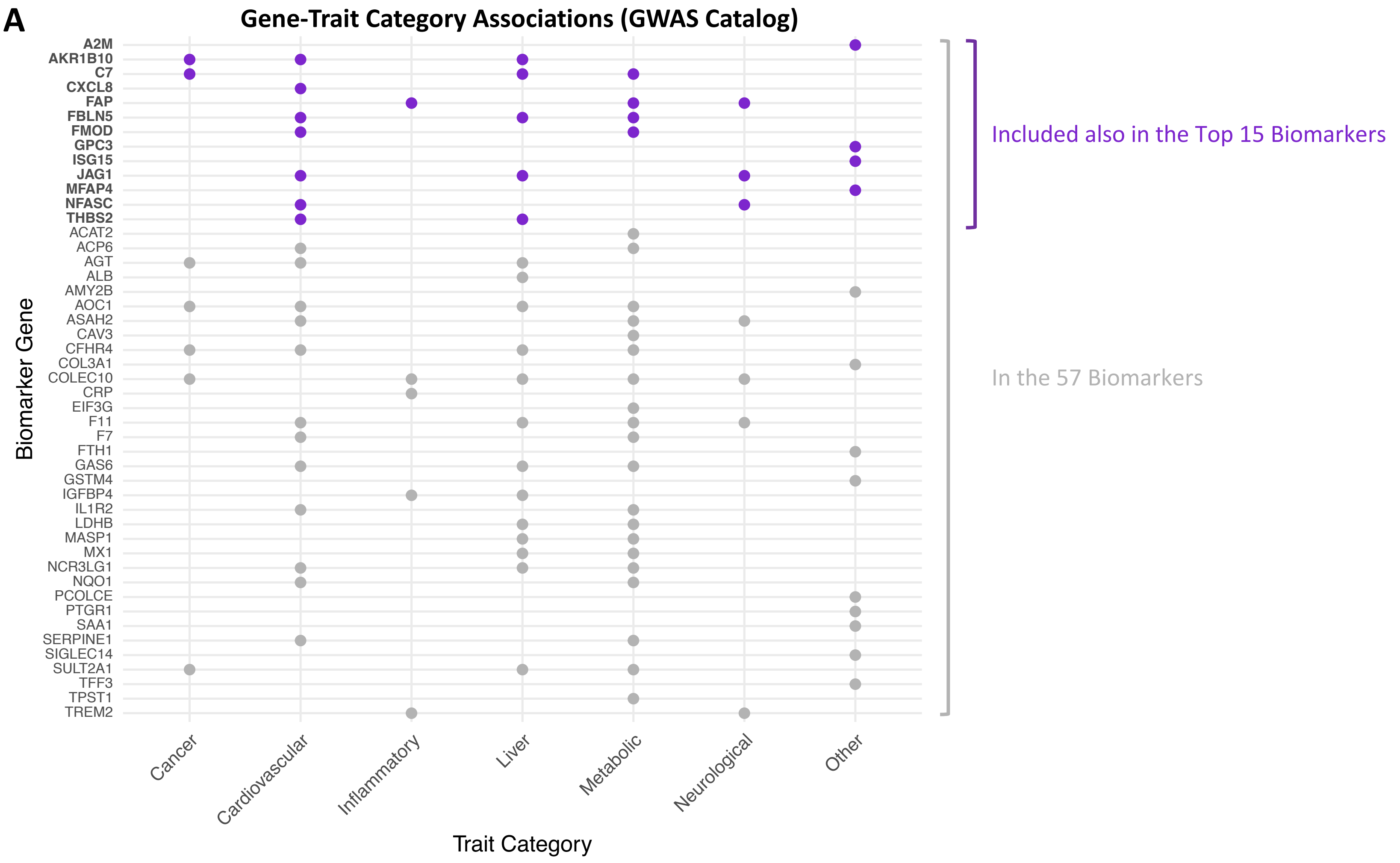

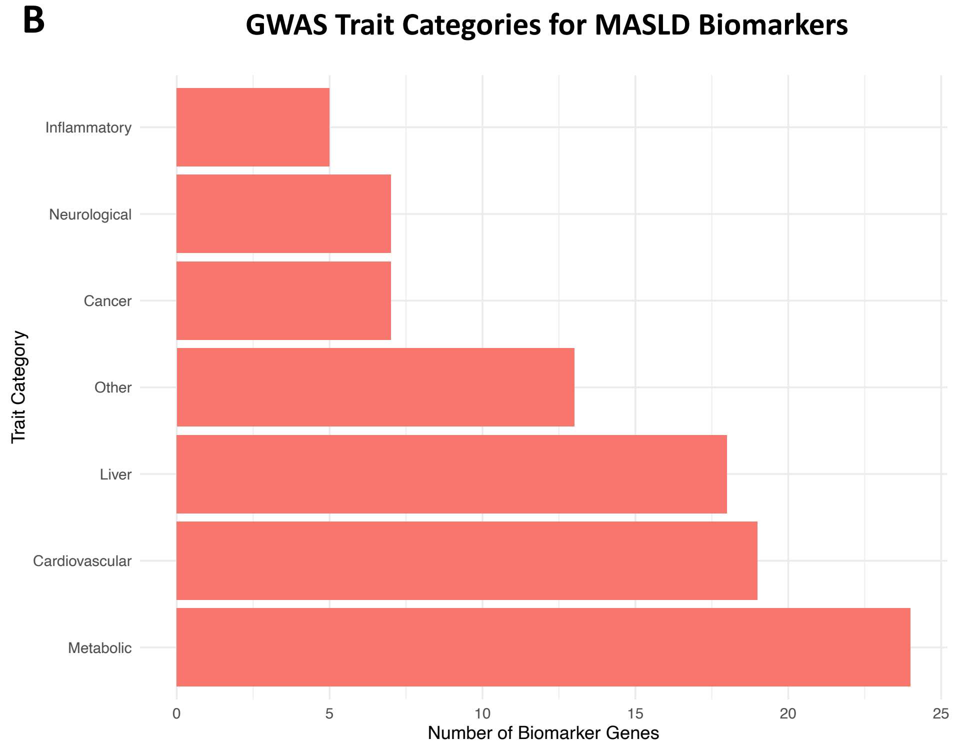

### Figure S11

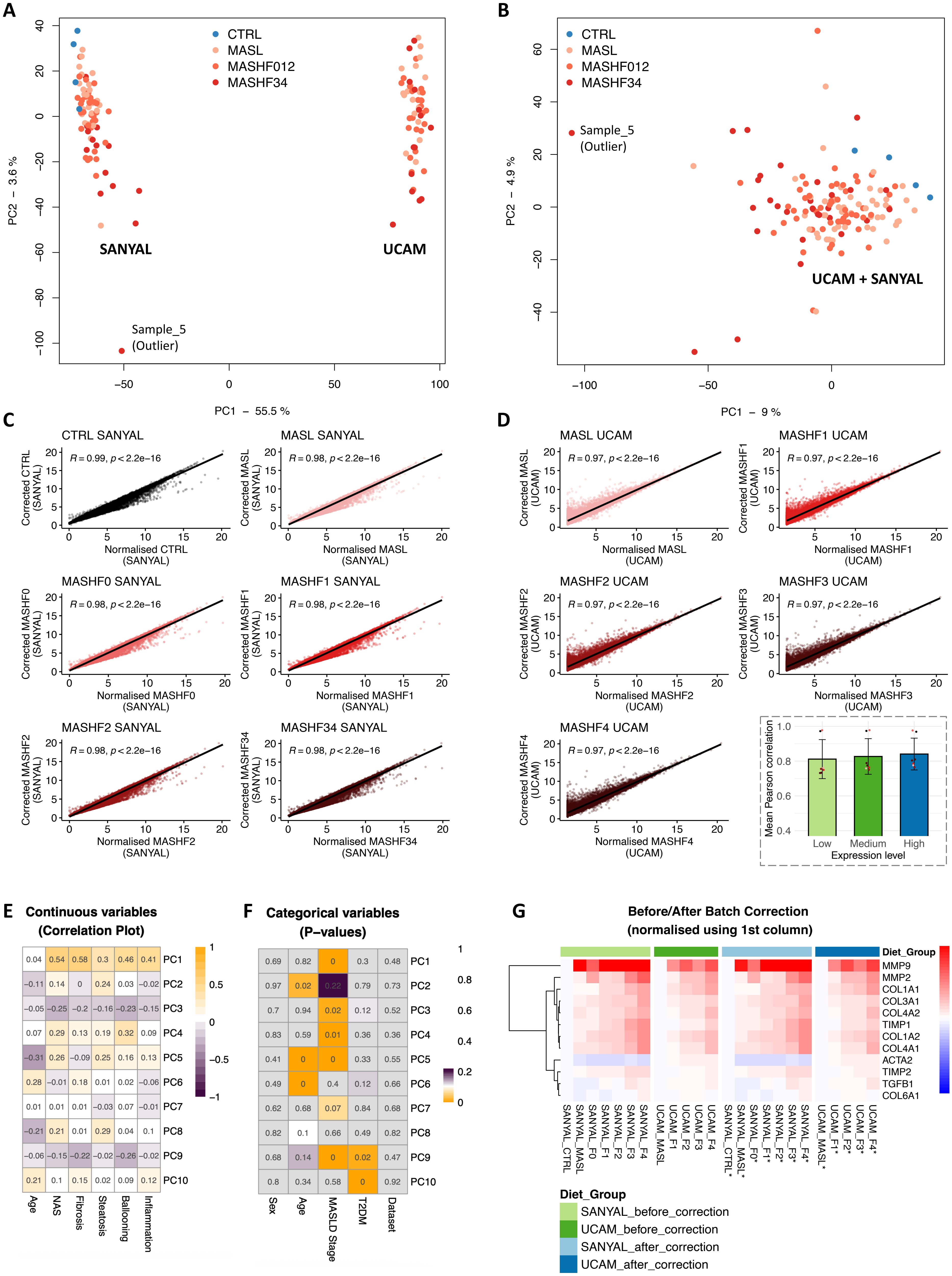
